# Adding Extra Knowledge in Scalable Learning of Sparse Differential Gaussian Graphical Models

**DOI:** 10.1101/716852

**Authors:** Arshdeep Sekhon, Beilun Wang, Yanjun Qi

## Abstract

We focus on integrating different types of extra knowledge (other than the observed samples) for estimating the sparse structure change between two *p*-dimensional Gaussian Graphical Models (i.e. differential GGMs). Previous differential GGM estimators either fail to include additional knowledge or cannot scale up to a high-dimensional (large *p*) situation. This paper proposes a novel method KDiffNet that incorporates Additional <u>K</u>nowledge in identifying <u>Diff</u>erential <u>Net</u>works via an Elementary Estimator. We design a novel hybrid norm as a superposition of two structured norms guided by the extra edge information and the additional node group knowledge. KDiffNet is solved through a fast parallel proximal algorithm, enabling it to work in large-scale settings. KDiffNet can incorporate various combinations of existing knowledge without re-designing the optimization. Through rigorous statistical analysis we show that, while considering more evidence, KDiffNet achieves the same convergence rate as the state-of-the-art. Empirically on multiple synthetic datasets and one real-world fMRI brain data, KDiffNet significantly outperforms the cutting edge baselines with regard to the prediction performance, while achieving the same level of time cost or less.

## 1 Introduction

Learning the change of dependencies between random variables is an essential task in many realworld applications. For example, when analyzing functional MRI samples from different groups of human subjects, detecting the difference in brain connectivity networks can shed light on studying and designing treatments for psychiatric diseases [6]. In this paper, we consider Gaussian graphical models (GGMs) and focus on directly estimating changes in the dependency structures of two *p*-dimensional GGMs, based on *n*_*c*_ and *n*_*d*_ samples drawn from the two models (we call the task differential GGMs). In particular, we focus on estimating the structure change under a high-dimensional situation, where the number of variables *p* may exceed the number of observations: *p >* max(*n*_*c*_, *n*_*d*_). To conduct consistent estimation under high dimensional settings, we leverage the sparsity constraint. In the context of estimating structural changes between two GGMs, this translates into a differential network with few edges. We review the state-of-the-art estimators for differential GGMs in Section 2.1.

One significant caveat of previous differential GGM estimators is that little attention has been paid to incorporating extra knowledge of the nodes or of the edges. In addition to the observed samples, extra information is widely available in real-world applications. For instance, when estimating the functional connectivity networks among brain regions via fMRI measurements (i.e. observed samples), there exist considerable knowledge about the spatial and anatomical evidence of these regions. Adding such evidence will help the learned differential structure better reflect domain experts’ knowledge like certain anatomical regions or spatially related regions are more likely to be connected [20].

Although being a strong evidence of structural patterns that we aim to discover, extra information has rarely been considered when estimating differential GGM from two sets of observed samples. To the authors’ best knowledge, only two loosely-related studies exist in the literature: (1) One study with the name NAK [2] (following ideas from [14]) proposed to integrate Additional Knowledge into the estimation of single-task graphical model using a weighted Neighbourhood selection formulation. (2) Another study with the name JEEK [18] (following [15]) considered edge-level evidence via a weighted objective formulation to estimate multiple dependency graphs from heterogeneous samples. Both studies only added edge-level extra knowledge in structural learning and neither of the approaches was designed for the direct structure estimation of differential GGM.

This paper fills the gap by proposing a novel method, namely KDiffNet, to add additional <u>K</u>nowledge in identifying <u>DIFF</u>erential networks via an Elementary Estimator. Our main objective is to make KDiffNet flexible enough to model various combinations of existing knowledge without re-designing the optimization. This is achieved by: (1) representing the edge-level domain knowledge as weights and using weights through a weighted 𝓁_1_ regularization constraint; and (2) describing the node-level knowledge as the variable groups and enforcing through a group norm constraint. Then KDiffNet designs a novel hybrid norm as the minimization objective and enforces the superposition of two aforementioned structured constraints. Our second main aim for KDiffNet is to achieve direct, scalable, and fast differential GGM estimations, and at the same time to guarantee the estimation error is well bounded. We achieve this goal through modeling KDiffNet in an elementary estimator based framework and solving it via parallel proximal based optimization. Briefly speaking, this paper makes the following contributions: ^1^

- **Novel and Flexible:** KDiffNet is the first method to integrate different kinds of additional knowledge for structure learning of differential GGMs. KDiffNet proposes a flexible formulation to consider both the edge-level evidence and the node-group level knowledge (Section 2.3).
- **Fast and Scalable:** We optimize KDiffNet through a proximal algorithm making it scalable to large values of *p*. KDiffNet ‘s unified formulation avoids the need to design knowledge-specific optimization (Section 2.5).
- **Theoretically Sound:** We theoretically prove the convergence rate of KDiffNet as 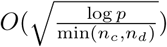, achieving the same error bound as the state-of-the-art (Section 2.6).
- **Empirical Evaluation:** We evaluate KDiffNet using multiple synthetic datasets and one realworld task. Our experiments showcase how KDiffNet can integrate knowledge of spatial distances, known edges or anatomical grouping evidence in the proposed formulation, empirically showing its real-world adaptivity. KDiffNet improves the state-of-the-art baselines with consistently better prediction accuracy while maintaining the same or less time cost (Section 3).

## 2 Proposed Method: KDiffNet

### 2.1 Previous Estimators for Structure Change between two GGMs (Differential GGMs)

The task of estimating differential GGMs assumes we are given two sets of observed samples (in the form of two matrices) 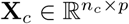 and 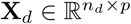, identically and independently drawn from two normal distributions *N*_*p*_(*µ*_*c*_, Σ_*c*_) and *N*_*p*_(*µ*_*d*_, Σ_*d*_) respectively. Here *µ*_*c*_, *µ*_*d*_ ∈ ℝ^*p*^ describe the mean vectors and Σ_*c*_, Σ_*d*_ ∈ ℝ^*p*×*p*^ represent covariance matrices. The goal of differential GGMs is to estimate the structural change Δ defined by [27] ^2^.

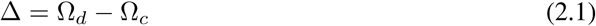

Here the precision matrices Ω_*c*_ := (Σ_*c*_)^−1^ and Ω_*d*_ := (Σ_*d*_)^−1^. The conditional dependency structure of a GGM is encoded by the sparsity pattern of its precision matrix. Therefore, one entry of Δ describes if the magnitude of conditional dependency of a pair of random variables changes between two conditions. A sparse Δ means few of its entries are non-zero, indicating a differential network with few edges.

A naive approach to estimate Δ is a two-step procedure in which we estimate 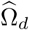 and 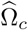 from two sets of samples separately and calculate 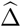 using Eq. (2.1). However, in a high-dimensional setting, the strategy needs to assume both Ω_*d*_ and Ω_*c*_ are sparse (to achieve consistent estimation of each), although the assumption is not necessarily true even if the change Δ is sparse (details in Section S:1).

Multiple recent studies have been motivated to directly estimate Δ from two sets of samples. We call these studies differential GGM estimators and group them to four kinds. (1) **Likelihood based**. Zhang et al. [25] used the fused norm for regularizing the maximum likelihood estimation (MLE) to simultaneously learn both two GGMs and the difference (*λ*_2_(‖Ω_*c*_‖_1_ + ‖Ω_*d*_‖_1_) + *λ*_*n*_‖Δ‖_1_). The resulting penalized MLE framework is a log-determinant program, which can be solved by block coordinate descent [25] or the alternating direction method of multipliers (ADMM) by the JGLFUSED package [5]. (2) **Density ratio based:**. Recently Liu et al. used density ratio estimation (SDRE) to directly learn Δ without having to identify the structures of Ω_*c*_ and Ω_*d*_. The authors focused on exponential family-based pairwise Markov networks [10] and solved the resulting optimization using proximal gradient descent [9]. (3) **Constrained** 𝓁_1_ **minimization based**. Diff-CLIME, another regularized convex program, was proposed to directly learn structural changes Δ without going through the learning of each individual GGMs [26]. It uses an 𝓁_1_ minimization formulation constrained by the covariance-precision matching, reducing the estimation problem to solving linear programs. All three aforementioned groups have used 𝓁_1_ regularized convex formulation for estimating Δ. (4) **Elementary estimator based**. The last category extends the so-called Elementary Estimator proposed by [21, 23, 22] to achieve a closed-form estimation of differential GGM via the DIFFEE estimator [19] (more in the next section and Section 2.4).

### 2.2 Background: Elementary Estimators for Graphical Models

#### 𝓁_1_ Regularized MLE for GGM Estimation: Graphical Lasso (GLasso)

The “GLasso” Estimator [24, 1] is the classic formulation for estimating sparse GGM from observations drawn from a single multivariate Gaussian distribution. It optimizes the following 𝓁_1_ penalized MLE objective:

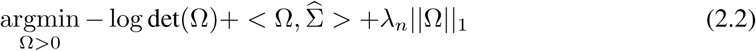

Where *λ*_*n*_ *>* 0 is the sparsity regularization parameter. While state-of-the-art optimization methods have been developed to solve the optimization in Eq. (2.2), they are expensive for large-scale tasks.

#### 𝓁_1_ based Elementary Estimator for Graphical Model (EE-GM) Estimation

Yang et al. [23] proposed to learn sparse Gaussian graphical model via the following formulation instead:

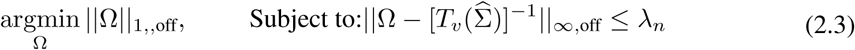

Actually [23] proposed the following generic formulation to estimate graphical models (GM) of exponential families (GGM is a special case of GM with exponential distribution):

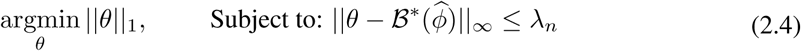

Here *θ* is the canonical parameter to be estimated and 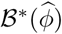 is a so-called proxy of backward mapping for the target GM. 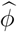 is the empirical mean of the sufficient statistics of the underlying exponential distribution. For example, in the case of Gaussian, *θ* is the precision matrix, 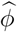 is the sample covariance matrix and the proxy backward mapping is 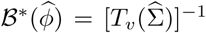 (We explain backward mapping, proxy backward mapping and the property and convergence rate of 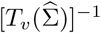 in Section S:4.1).

The main advantage of Eq. (2.4) and Eq. (2.3) was that they are simple estimators with computationally easy solutions. Importantly their solutions achieve the same sharp convergence rate as the regularized convex formulation of Eq. (2.2) when under high-dimensional settings.

#### ℛ(·) norm based Elementary Estimators

Recently multiple studies [21, 22, 19, 18] followed [23] and expanded EE-GM into a more general framework “Elementary estimators” (EE):

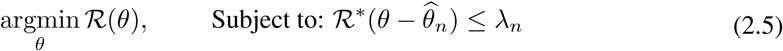

Where ℛ (·) represents a decomposable regularization function. ℛ^*^(·) is the dual norm of ℛ(·),

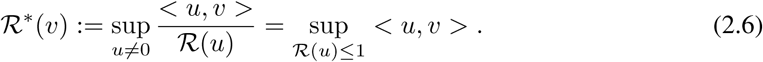

Eq. (2.4) and Eq. (2.3) are special cases of Eq. (2.5). 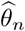 needs to be carefully constructed, well-defined and closed-form for the purpose of simplified computations. For example, [21] conduct the high-dimensional estimation of linear regression models by using the classical ridge estimator as 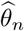 in Eq. (2.5). When 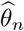 itself is closed-form and comes with strong statistical convergence guarantees in high-dimensional situations, we can use the unified framework proposed by the recent seminal study from [11] to prove that the solution of Eq. (2.5) achieves the near optimal convergence rate as comparable to regularized convex formulations when satisfying certain conditions.

### 2.3 Integrating additional knowledge and Δ with a Novel Function: kEV norm

Section 2.1 points out that none of the previous Δ estimators have designed to integrate extra evidence beyond two sets of observed samples. Differently our Δ estimator aims to achieve two goals: (1) the new estimator should be flexible enough to describe various kinds of real-world knowledge, including like spatial distance, hub knowledge, known interactions or how multiple variables function as groups (see below). (2) the new estimator should work well in high-dimensional situations (large *p*) and is computationally practical. Eq. (2.5) provides an intriguing formulation to build simpler and possibly fast estimators accompanied by statistical guarantees, as long as 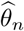 can be carefully constructed, well-defined and closed-form. We adapt it to design KDiffNet in the next Section 2.4.

In order to use Eq. (2.5) for estimating our target parameter *θ* = Δ, we need to design ℛ (Δ).

#### (1) Knowledge as Weight Matrix

We can describe the edge-level knowledge as positive weight matrices like *W*_*E*_ ∈ ℝ^*p*×*p*^. For example, when estimating the functional brain connectivity networks among brain regions *W*_*E*_ can describe spatial distance among brain regions that are publicly available through projects like openfMRI [12]. Another important example is when identifying gene-gene interactions from patients’ gene expression profiles. Besides the patient samples, state-of-the-art bio-databases like HPRD [13] have collected a significant amount of information about direct physical interactions among corresponding proteins, regulatory gene pairs or signaling relationships collected from high-quality bio-experiments. Here *W*_*E*_ can describe existing known edges as the knowledge, like those from interaction databases for discovering gene networks (a semi-supervised setting for such sample based network estimations).

The positive matrix-based representation provides a powerful and flexible strategy that allows integration of many possible forms of existing knowledge to improve differential structure estimation, as long as they can be represented into edge-level weights. We can combine *W*_*E*_ knowledge and the sparse regularization of Δ into a weighted 𝓁_1_ norm ‖*W*_*E*_ ∘ Δ‖_1_, enforcing prior known importance of edges in the differential graph through weights. The larger a weight entry in *W*_*E*_, the less likely the corresponding edge belongs to the true differential graph. As mentioned in Section 1, NAK and JEEK estimators have tried similar weight matrix based strategy to add extra knowledge in identifying single-task GGM and in discovering multiple GGMs. None of the previous differential GGM estimators have explored this though.

#### (2) Knowledge as Node Groups

In many real-world applications, there exist known group knowledge about random variables. For example, when working with genomics samples, biologists have collected a rich set of group evidence about how genes belong to various biological pathways or exist in the same or different cellular locations [4]. Such knowledge of node grouping provides solid biological bias like genes belonging to the same biology pathway tend to have interactions among them (shared dependency pattern) in one cellular context or tend to not interact with each other (shared sparsity) at some other cellular conditions. However, this type of group evidence cannot be described via the weight matrix *W*_*E*_ based formulation.

This is because even though it is safe to assume nodes in the same group share similar interaction patterns, but we do not know beforehand if the nodes in the group are collectively part of the differential network (group dependency) or not (group sparsity). To mitigate this issue, we use a flexible known node-group norm to include such extra knowledge. We represent the group knowledge as a set of groups on feature variables (vertices) 𝒢_*p*_. Formally, *∀g*_*k*_ ∈ 𝒢_*p*_, *g*_*k*_ = {*i*} where *i* indicates that the *i*-th node belongs to the group *k*. Integrating 𝒢_*p*_ knowledge into Δ means to enforce a group sparsity regularization on Δ. We generate edge-group index 𝒢_*V*_ from the node group index 𝒢_*p*_. This is done via defining 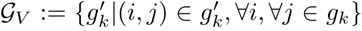. For vertex nodes in each node group *g*_*k*_, all possible pairs between these nodes belong to an edge-group 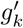. We propose to use the group,2 norm 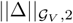 to enforce group-wise sparse structure on Δ. None of the previous differential GGM estimators have explored this knowledge-integration strategy before.

#### kEV norm

Now we propose a novel norm ℛ(Δ) to combine the two strategies above. We assume that the true parameter 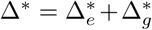 is a superposition of two “clean” structures, a sparse structured 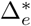 and a group-structured 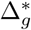. We propose a new norm, <u>k</u>nowledge for <u>E</u>dges and <u>V</u>ertex norm (kEV-norm), as the superposition of the edge-weighted 𝓁_1_ norm and the group structured norm:

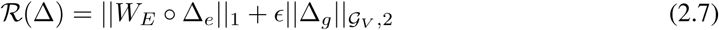

Our target parameter Δ = Δ_*e*_ + Δ_*g*_. The Hadamard product ∘ is the element-wise product between two matrices i.e. [*A* ∘ *B*]_*ij*_ = *A*_*ij*_*B*_*ij*_ and 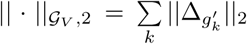 where *k* is the *k*-th group.

*W*_*E*_ ∈ *R*^*p*×*p*^ is the aforementioned edge-level additional knowledge. *W*_*E*_ *>* 0, *∀ i, j* ∈ {1 … *p*}. *ϵ >* 0 is a hyperparameter. kEV-norm has the following three properties (proofs in Section S:3).

- (i) kEV-norm is a norm function if *ϵ* and entries of *W*_*E*_ are positive.
- (ii)If the condition in (i) holds, kEV-norm is a decomposable norm.
- (iii)The dual norm of kEV-norm is

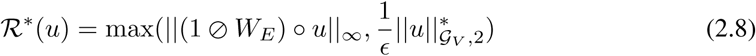

### 2.4 KDiffNet : <u>k</u>EV Norm based Elementary Estimator for identifying <u>Diff</u>erential <u>Net</u>

Our goal is to achieve simple, scalable and theoretically sound estimation. EE in Eq. (2.5) provides such a formulation as long as we can construct 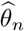 well. Now we have ℛ(Δ) as Eq. (2.7) and its dual norm ℛ ^*^(·) in Eq. (2.8). We just need to find 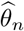 for Δ that is carefully constructed, theoretically well-behaved when high-dimensional, and closed-form for the purpose of simplified computations.

One key insight of differential GGM is that the density ratio of two Gaussian distributions is naturally an exponential-family distribution (see proofs in Section S:4.2). The differential network Δ is one entry of the canonical parameter for this distribution. The MLE solution of estimating vanilla (i.e. no sparsity and not high-dimensional) graphical model in an exponential family distribution can be expressed as a backward mapping that computes the target model parameters from certain given moments. When using vanilla MLE to learn the exponential distribution about differential GGM (i.e., estimating canonical parameter), the backward mapping of Δ can be easily inferred from the two sample covariance matrices using 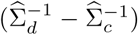 (Section S:4.2). Even though this backward mapping has a simple closed form, it is not well-defined when high-dimensional because 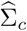 and 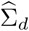 are rank-deficient (thus not invertible) when *p > n*. Using Eq. (2.3) to estimate Δ, Wang et. al. [19] proposed the DIFFEE estimator for EE-based differential GGM estimation and used only the sparsity assumption on Δ. This study proposed a proxy backward mapping as 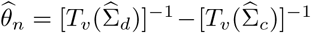. Here [*T*_*v*_(*A*)]_*ij*_ := *ρ*_*v*_(*A*_*ij*_) and *ρ*_*v*_(·) is chosen as a soft-threshold function.

We borrow the idea to use 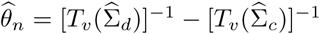. In Section S:4.3 and Section S:4.4 we prove that 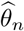 is both available in closed-form, and well-defined in high-dimensional settings. Now by plugging ℛ(Δ), its dual ℛ^*^(·) and 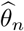 into Eq. (2.5), we get the formulation of KDiffNet :

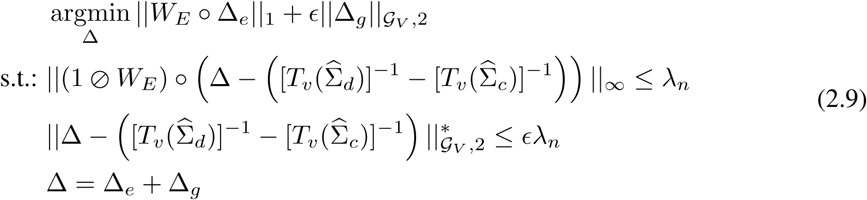

### 2.5 Solving KDiffNet

We then propose to use a proximal parallel based optimization to solve Eq. (2.9), inspired by its distributed and parallel nature [3]. To simplify notations, we add a new notation Δ_*tot*_ := [Δ_*e*_; Δ_*g*_], where; denotes the row wise concatenation. We also add three operator notations including *L*_*e*_(Δ_*tot*_) = Δ_*e*_, *L*_*g*_(Δ_*tot*_) = Δ_*g*_ and *L*_*tot*_(Δ_*tot*_) = Δ_*e*_ + Δ_*g*_. Now we obtain the following re-formulation of KDiffNet :

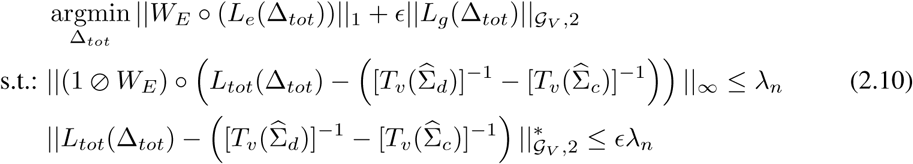

Actually the three added operators are affine mappings: *L*_*e*_ = *A*_*e*_Δ_*tot*_, *L*_*g*_ = *A*_*g*_Δ_*tot*_, and *L*_*tot*_ = *A*_*tot*_Δ_*tot*_, where *A*_*e*_ = [***I***_*p*×*p*_ **0**_*p*×*p*_], *A*_*g*_ = [**0**_*p*×*p*_ ***I***_*p*×*p*_] and *A*_*tot*_ = [***I***_*p*×*p*_ ***I***_*p*×*p*_].

Algorithm 1 summarizes the Parallel Proximal algorithm [3, 22] we propose for optimizing Eq. (2.10). In Section S:1.3 we further prove that its computational cost is *O*(*p*^3^). More concretely in Algorithm 1, we simplify the notations by denoting 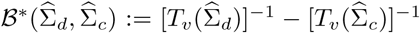, and reformulate Eq. (2.10) to the following equivalent and distributed formulation:

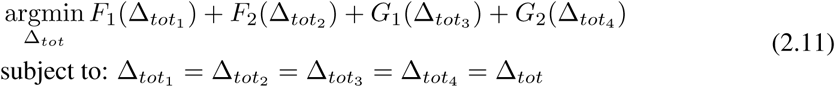

Where 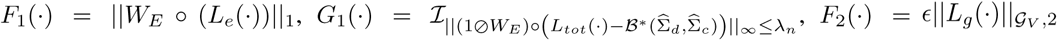 and 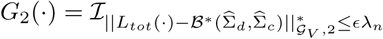. Here ℐ_*C*_ (·) represents the indicator function of a convex set *C* denoting that ℐ_*C*_ (*x*) = 0 when *x* ∈ *C* and otherwise ℐ_*C*_ (*x*) = *∞*. The detailed solution of each proximal operator is summarized in Table S:1 and Section S:2.

#### Algorithm 1

A Parallel Proximal Algorithm to optimize KDiffNet

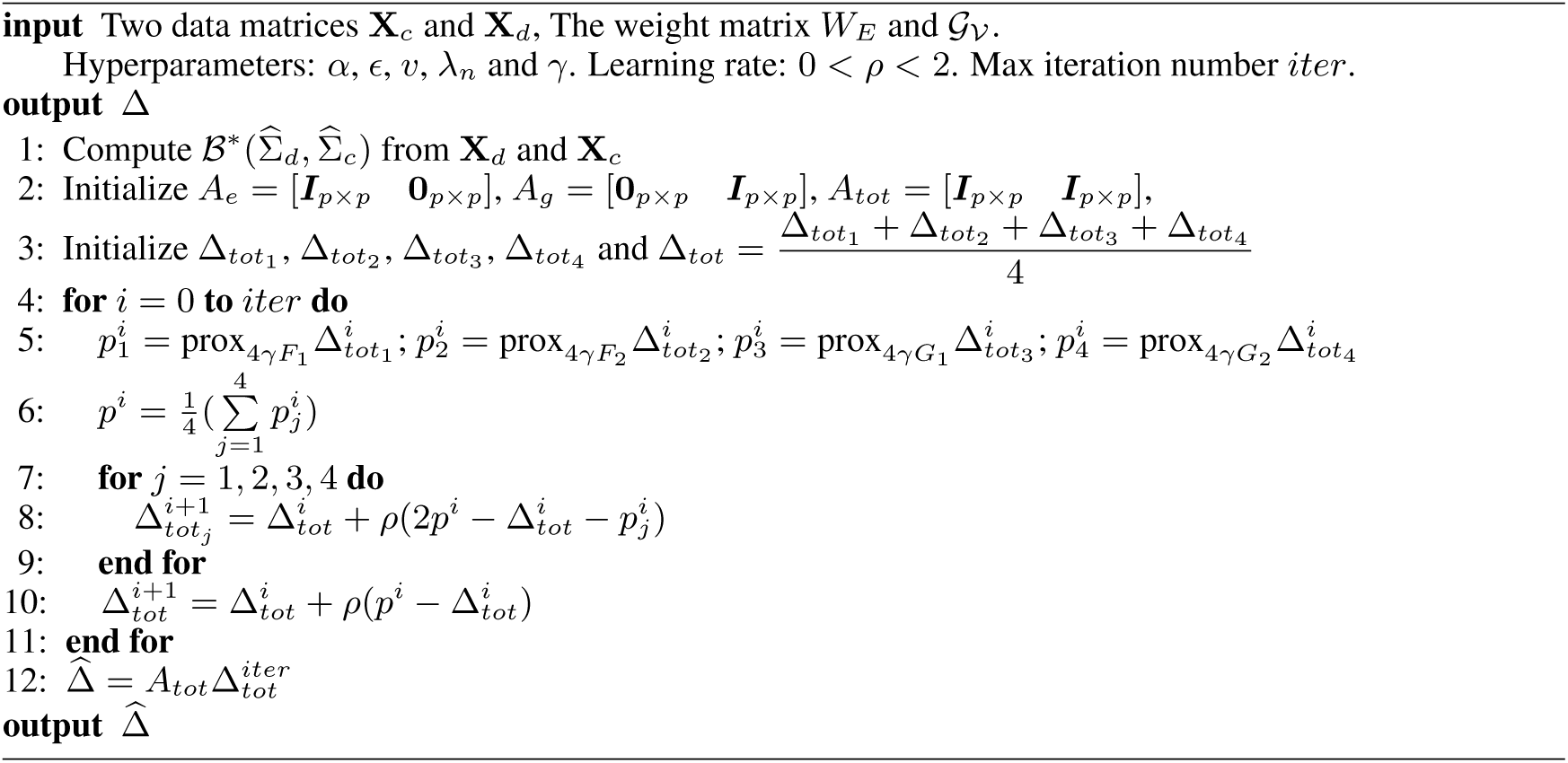

### 2.6 Analysis of Error Bounds

Based on Theorem S:5.3 and conditions in Section S:5, we have the following corollary about the convergence rate of KDiffNet. See its proof in Section S:5.2.2.

#### Corollary 2.1.

*In the high-dimensional setting, i.e., p >* max(*n*_*c*_, *n*_*d*_), *let* 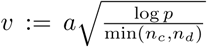. *Then for* 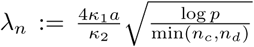 *and* min(*n*_*c*_, *n*_*d*_) *> c* log *p, with a probability of at least* 1 −2*C*_1_ exp(*-C*_2_*p* log(*p*)), *the estimated optimal solution* 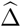 *has the following error bound:*

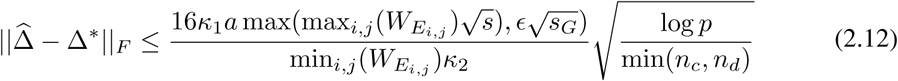

*where C*_1_,*C*_2_,*a, c, κ*_1_ *and κ*_2_ *are constants. See s and s*_*G*_ *in Definition S:3.4.*

## 3 Experiments

We aim to empirically show that KDiffNet is adaptive and flexible in incorporating different kinds of available evidence for improved differential network estimation. **Data:** This is accomplished by evaluating KDiffNet and baselines on two sets of datasets: (1) A total of 126 different synthetic datasets representing various combinations of additional knowledge (details see Section 3.1); and (2) one real-world fMRI dataset ABIDE for functional brain connectivity estimation (Section 3.2). We obtain the edge-level knowledge from three different human brain atlas [7, 8, 16] about brain connectivity, resulting in three different *W*_*E*_ with *p* = {116, 160, 246}. For each atlas we compute *W*_*E*_ using the spatial distance between its brain Region of Interests (ROIs). At the same time, we explore two different types of group knowledge about brain regions from Dosenbach Atlas[7] (Section 3.2). **Baselines:** We compare KDiffNet to JEEK[18] and NAK[2], that use the extra edge knowledge, two direct differential estimators (SDRE[9], DIFFEE[19]) and MLE based JGLFUSED[5] (Section 2.1 and detailed equations of each in Section S:1). We also extend KDiffNet to data situations with only edge knowledge (KDiffNet-E) or only group knowledge (KDiffNet-G). Both variations (KDiffNet-E and KDiffNet-G) can be solved by fast closed form solutions (Section S:2.2).

Additional details of setup, metrics and hyper-parameters are in Section S:6.1. **Hyperparameters:** The key hyper-parameters are tuned as follows:

- *v* : To compute the proxy backward mapping, we vary *v* in {0.001*i*|*i* = 1, 2, …, 1000} (to make *T*_*v*_(Σ_*c*_) and *T*_*v*_(Σ_*d*_) invertible).
- *λ*_*n*_ : According to our convergence rate analysis in Section 2.6, 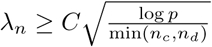, we choose *λ*_*n*_ from a range of 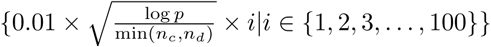 using cross-validation. For KDiffNet-G, we tune over *λ*_*n*_ from a range of 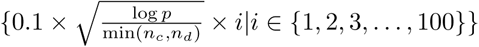^3^.
- *ϵ*: For KDiffNet-EG experiments, we tune *ϵ* ∈ {0.0001, 0.01, 1, 100}.

### 3.1 Experiment: Simulated Data about Brain Connectivity using Three Real-World Brain Spatial Matrices and Anatomic Group Evidence from Neuroscience as Knowledge

In this section, we show the effectiveness of KDiffNet in integrating additional evidence through a comprehensive set of many simulation datasets. Our simulated data settings mimic three possible types of additional knowledge in the real-world: with both edge and known node group knowledge (Data-EG), with only edge-level evidence (Data-E) or with only known node groups (Data-G). For the edge knowledge, we consider three cases of *W*_*E*_ with *p* = {116, 160, 246} computed from three human brain atlas about brain regions [7, 8, 16]. For the group knowledge, we simulate groups to represent related anatomic regions inspired by the atlas [7]. For each simulation dataset, two blocks of data samples are generated following Gaussian distribution using 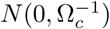 and 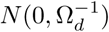 via the simulated Ω_*c*_ and Ω_*d*_. Each simulated dataset includes a pair of data blocks to estimate its differential GGM. We conduct a comprehensive evaluation over a total of 126 different simulated datasets by varying (*p*), varying the number of samples (*n*_*c*_ and *n*_*d*_), changing the proportion of edges controlled by *W*_*E*_ (*s*) and by varying the number of known groups *s*_*G*_. The details of the simulation framework are in Section S:6.2.

We present a summary of our results (partial) in Table 1 using columns showing two cases of data generation settings (Data-EG and Data-G). Table 1 uses the mean F1-score and the computational time cost to compare methods (rows). Results about simulated datasets under Data-E case are in Section S:6. We can make several conclusions:

**Table 1:** Mean Performance(F1-Score) and Computation Time(seconds) with standard deviation given in parentheses for the same setting of *n*_*c*_ and *n*_*d*_ of KDiffNet-EG, KDiffNet-E, KDiffNet-G and baselines for simulated data. * indicates that the method is not applicable for a data setting.

| Method | Data-EG(Time) | Data-EG(F1-Score) |  |  | Data-G(Time) | Data-G(F1-Score) |
| --- | --- | --- | --- | --- | --- | --- |
| | W2($p = 246$) | W1($p = 116$) | W2($p = 246$) | W3($p = 160$) | W2($p = 246$) | W2($p = 246$) |
| KDiffNet-EG | 4.591±0.08 | **0.717±0.05** | **0.927±0.01** | **0.934±0.07** | * | * |
| KDiffNet-G | **0.013±0.01** | 0.578±0.09 | 0.565±0.10 | 0.575±0.09 | 0.006±0.00 | **0.891±0.06** |
| KDiffNet-E | **0.017±0.01** | 0.692±0.06 | 0.872±0.02 | 0.916±0.02 | * | * |
| JEEK [18] | 10.998±0.11 | 0.572±0.10 | 0.581±0.09 | 0.580±0.09 | * | * |
| NAK[2] | 3.800±0.15 | 0.226±0.08 | 0.204±0.07 | 0.207±0.08 | * | * |
| SDRE[10] | 27.487±2.59 | 0.573±0.11 | 0.568±0.11 | 0.574±0.11 | 11.764±1.23 | 0.318±0.10 |
| DIFEE[19] | **0.004±0.00** | 0.570±0.11 | 0.560±0.11 | 0.570±0.11 | 0.004±0.00 | 0.135±0.02 |
| JGLFUSED[5] | 68.128±1.66 | 0.502±0.08 | 0.481±0.08 | 0.495±0.08 | 144.470±59.68 | 0.055±0.01 |

1. **KDiffNet outperforms those baselines not considering knowledge.** Clearly KDiffNet and its variations achieve the highest F1-score across all the 126 datasets. SDRE and DIFFEE are direct differential network estimators but perform poorly indicating that adding additional knowledge improves differential GGM estimation. MLE based JGLFUSED performs the worst in all cases.
2. **KDiffNet outperforms those baselines considering knowledge, especially when group knowledge exist.** When under the Data-EG setting, while JEEK and NAK include the extra edge information, they cannot integrate group information and are not for differential estimation. This results in lower F1-Score(0.581 and 0.204 for W2) compared to KDiffNet-EG (0.927 for W2). The advantage of modeling both edge and node groups evidence is also indicated by the higher F1-Score of KDiffNet-EG with respect to KDiffNet-E and KDiffNet-G on the Data-EG setting (Top 3 rows in Table 1). On Data-G cases, none of the baselines can model node group evidence. On average KDiffNet-G performs 6.4× better than the baselines for *p* = 246 with respect to F1.
3. **KDiffNet achieves reasonable time cost versus the baselines and is scalable to large** *p*. Figure 1(a) shows each method’s time cost per *λ*_*n*_ for large *p* = 2000. Consistently KDiffNet-EG is faster than JEEK, JGLFUSED and SDRE (Column 1 in Table 1). KDiffNet-E and KDiffNet-G are faster than KDiffNet-EG owing to closed form solutions. On Data-G dataset and Data-E datasets (Section S:6.2), our faster closed form solutions achieve much more significant computational all the baselines. For example on datasets using W2 *p* = 246, KDiffNet-E and KDiffNet-G are on an average 21000× and 7400× faster (Column 5 in Table 1) than the baselines, respectively. We have all detailed results and figures about F1-Score and time cost for all 126 data settings in Section S:6.2. Besides F1-Score, we also present the ROC curves from all methods when varying *λ*_*n*_. KDiffNet achieves the highest Area under Curve (AUC) in comparison to all other baselines.

### 3.2 Experiment: Functional Connectivity Estimation from Real-World Brain fMRI Data

In this experiment, we evaluate KDiffNet in a real-world downstream classification task on a publicly available resting-state fMRI dataset: ABIDE[6]. This aims to understand how functional dependencies among brain regions vary between normal and abnormal and help to discover contributing markers that influence or cause the neural disorders [17]. ABIDE includes two groups of human subjects: autism and control. We utilize three types of additional knowledge: *W*_*E*_ based on the spatial distance between 160 regions of the brain[7] and two types of available node groups from Dosenbach Atlas[7]: one with 40 unique groups about macroscopic brain structures (G1) and another with 6 higher level node groups having the same functional connectivity(G2). We use Quadratic Discriminant Analysis (QDA) in downstream classification to assess the ability of the estimators to learn the differential patterns about the connectome structures. (Details of the ABIDE dataset, baselines, design of the additional knowledge *W*_*E*_ matrix, cross-validation and the QDA classification method are in Section S:6.4.) Figure 1(b) compares KDiffNet-EG, KDiffNet-E, KDiffNet-G and baselines on ABIDE, using the *y* axis for classification test accuracy (the higher the better) and the *x* axis for the computation speed (negative log seconds, the more right the better). KDiffNet-EG1, incorporating both edge(*W*_*E*_) and (G1) group knowledge, achieves the highest accuracy of 57.2% for distinguishing the autism subjects versus the control subjects without sacrificing computation speed ^4^.

**Figure 1:**
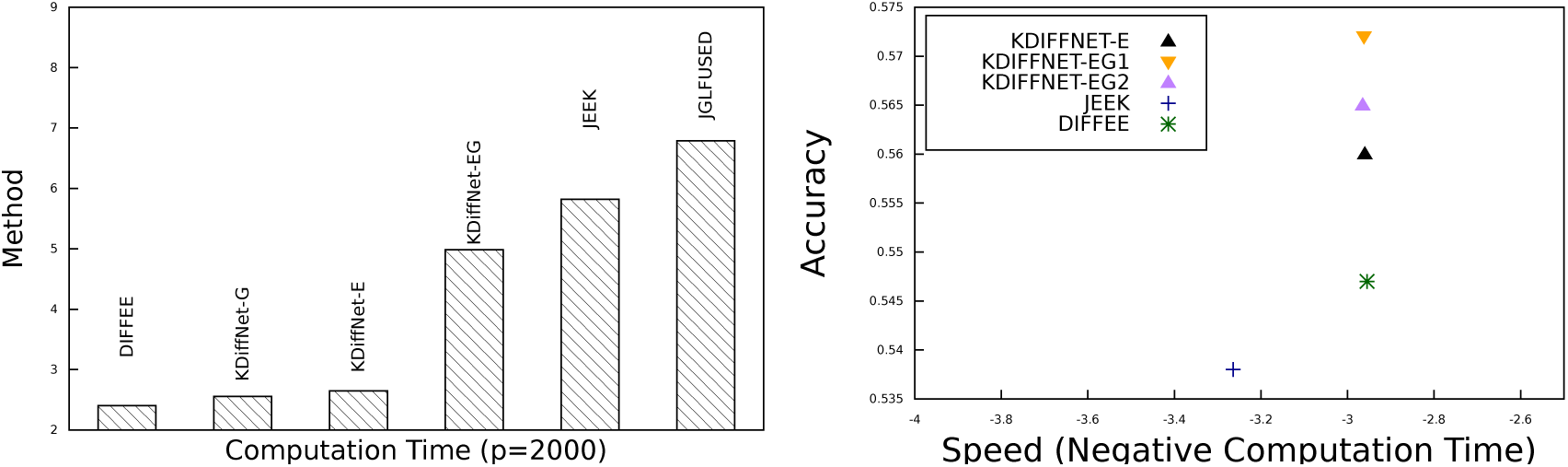
(a)(LEFT) Computation Time (log milliseconds) per *λ*_*n*_ for large *p* = 2000: KDiffNet-EG has reasonable time cost with respect to baseline methods. KDiffNet-E and KDiffNet-G are fast closed form solutions. (b) (RIGHT) ABIDE Dataset: KDiffNet-EG achieves highest Accuracy without sacrificing computation speed (points towards top right are better).

## 4 Conclusions

We propose a novel method, KDiffNet, to incorporate additional knowledge in estimating differential GGMs. KDiffNet elegantly formulates existing knowledge based on the problem at hand and avoids the need to design knowledge-specific optimization. We sincerely believe the scalability and flexibility provided by KDiffNet can make differential structure learning of GGMs feasible in many real-world tasks. We plan to generalize KDiffNet from Gaussian to semi-parametric distributions or to Ising Model structures. As node group knowledge is particularly important and abundant in genomics, we plan to evaluate KDiffNet on more real-world genomics data with multiple types of group information.

## Acknowledgement

This work was supported partly by the National Science Foundation under NSF CAREER award No. 1453580. Any Opinions, findings and conclusions or recommendations expressed in this material are those of the author(s) and do not necessarily reflect those of the National Science Foundation.

### Appendix

#### S:1 Connecting to Relevant Studies

##### S:1.1 Differential GGM Estimation

**JGLFUSED[6]:** This study extends the previously mentioned MLE based GLasso(Section 2.2) estimator for sparse differential GGM estimation. An additional sparsity penalty on the differential network, called the fused norm, is included as part of the optimization objective:

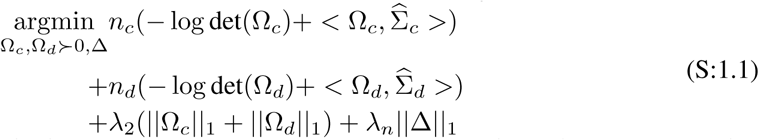

The alternating direction method of multipliers (ADMM) method was used to solve Eq. (S:1.1) that needs to run expensive SVD in one sub-procedure [6].

**DIFFEE[23]:** Computationally, EEs are much faster than their regularized convex program peers for GM estimation. [23] proposed the so-called DIFFEE for estimating sparse changes in high-dimensional GGM structure using EE:

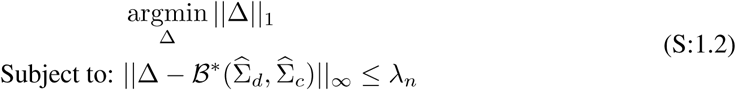

[23] use a closed form and well-defined proxy function 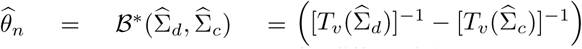 to approximate the backward mapping (the vanilla MLE solution) for differential sGGMs. We explain the proxy backward mapping and its statistical properties in Section S:4.1. The DIFFEE solution is a closed-form entry-wise thresholding operation on 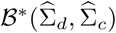 to ensure the desired sparsity structure of its final estimate. Here *λ*_*n*_ *>* 0 is the tuning parameter. Eq. (S:1.2) is a special case of Eq. (2.5), in which ℛ(·) is the 𝓁 _1_-norm for sparsity and the differential network Δ is the *θ* we aim to estimate.

As claimed by [10] direct estimation of differential GGMs can be more efficient both in terms of the number of required samples as well as the computation time cost. Besides, it does not require to assume each precision matrix as sparse. For instance recent literature in neuroscience has suggested that each subject’s functional brain connection network may not be sparse, even though differences across subjects may be sparse [1]. When identifying how genetic networks vary between two conditions, each individual network may contain hub nodes, therefore not entirely sparse [11].

**SDRE[12]:** [12] proposed to estimate Sparse differential networks in exponential families by Density <u>R</u>atio Estimation using the following formulation:

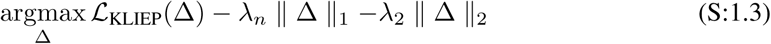

ℒ _KLIEP_ minimizes the KL divergence between the true probability density *p*_*d*_(*x*) and the estimated 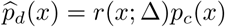 without explicitly modeling the true *p*_*c*_(*x*) and *p*_*d*_(*x*). This estimator uses the elastic-net penalty for enforcing sparsity. We use the sparseKLIEP^1^, that uses sub-gradient descent optimization as a baseline to our method.

**Diff-CLIME[25]:** This study directly learns Δ through a constrained optimization formulation.

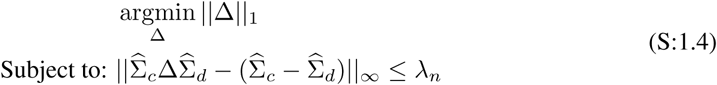

The optimization reduces to multiple linear programming problems, which in turn makes this method less scalable to large *p* with a computational complexity of *O*(*p*^8^).

##### S:1.2 Incorporating Additional Knowledge in GGM Estimation

While previous studies do not use available additional knowledge for differential structure estimation, a few studies have tried to incorporate edge level weights for other types of GGM estimation.

**NAK [3]:** For the single task sGGM, one recent study [3] (following ideas from [17]) proposed to use a weighted <u>N</u>eighborhood selection formulation to integrate edge-level <u>A</u>dditional <u>K</u>nowledge (NAK) as: 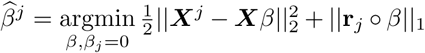. Here 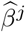 is the *j*-th column of a single sGGM 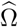. Specifically, 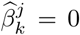 if and only if 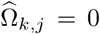. **r**_*j*_ represents a weight vector designed using available extra knowledge for estimating a brain connectivity network from samples ***X*** drawn from a single condition. The NAK formulation can be solved by a classic Lasso solver like glmnet.

**JEEK[22]:** Two related studies, JEEK[22] and W-SIMULE[18] incorporate edge-level extra knowledge in the joint discovery of *K* heterogeneous graphs. In both these studies, each sGGM corresponding to a condition *i* is assumed to be composed of a task specific sGGM component 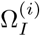 and a shared component Ω_*S*_ across all conditions, i.e., 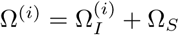. The minimization objective of W-SIMULE is as follows: objective:

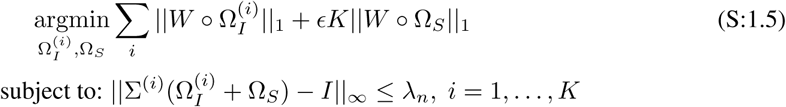

W-SIMULE is very slow when *p >* 200 due to the expensive computation cost *O*(*K*^4^*p*^5^). In comparison, JEEK is an EE-based optimization formulation:

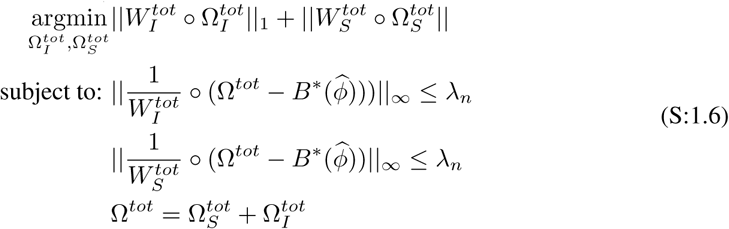

Here, 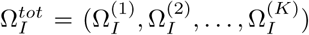 and 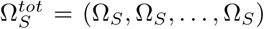. The edge knowledge of the task-specific graph is represented as weight matrix *W* ^(*i*)^ and *W*_*S*_ for the shared network. JEEK differs from W-SIMULE in its constraint formulation, that in turn makes its optimization much faster and scalable than WSIMULE. In our experiments, we use JEEK as our baseline.

**Drawbacks:** However, none of these studies are flexible to incorporate other types of additional knowledge like node groups or cases where overlapping group and edge knowledge are available for the same target parameter. Further, these studies are limited by the assumption of sparse single condition graphs. Estimating a sparse difference graph directly is more flexible as it does not rely on this assumption.

##### S:1.3 Computational Complexity

We optimize KDiffNet through a proximal algorithm, while KDiffNet-E and KDiffNet-G through closed-form solutions. The resulting computational cost for KDiffNet is *O*(*p*^3^), broken down into the following steps:

- Estimating two covariance matrices: The computational complexity is *O*(*max*(*n*_*c*_, *n*_*d*_)*p*^2^).
- Backward Mapping: The element-wise soft-thresholding operation [*T*_*v*_(·)] on the estimated covariance matrices, that costs *O*(*p*^2^). This is followed by matrix inversions [*T*_*v*_(·)]^−1^ to get the proxy backward mapping, that cost *O*(*p*^3^).
- Optimization: For KDiffNet, each operation in the proximal algorithm is group entry wise or entry wise, the resulting computational cost is *O*(*p*^2^). In addition, the matrix multiplications cost *O*(*p*^3^).

For KDiffNet-E and KDiffNet-G versions, the solution is the element-wise soft-thresholding operation 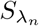, that costs *O*(*p*^2^).

**Figure S:1:**
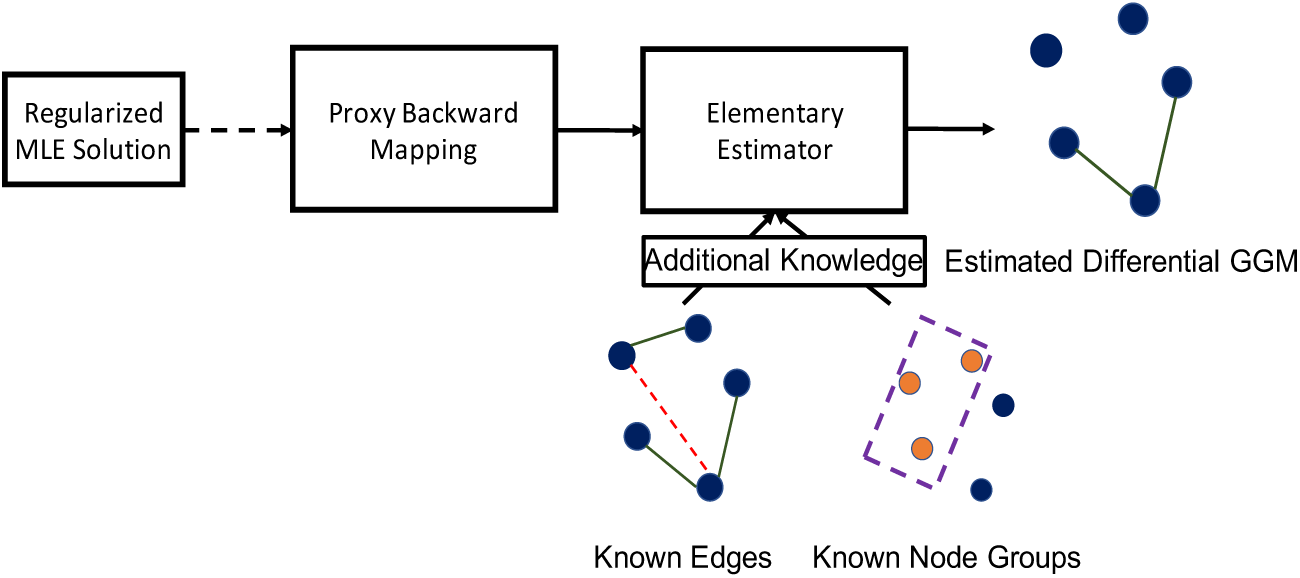
Schematic Diagram of KDiffNet : integrating extra edge and node groups knowledge for directly estimating the sparse change in the dependency structures of two *p*-dimensional GGMs (differential GGMs)

#### S:2 Optimization of KDiffNet and Its Variants

##### S:2.1 Optimization via Proximal Solution

We assume Δ_*tot*_ = [Δ_*e*_; Δ_*g*_], where; denotes row wise concatenation. Consider operator *L*_*d*_(Δ_*tot*_) = Δ_*e*_ and *L*_*g*_(Δ_*tot*_) = Δ_*g*_, *L*_*tot*_(Δ_*tot*_) = Δ_*e*_ + Δ_*g*_.

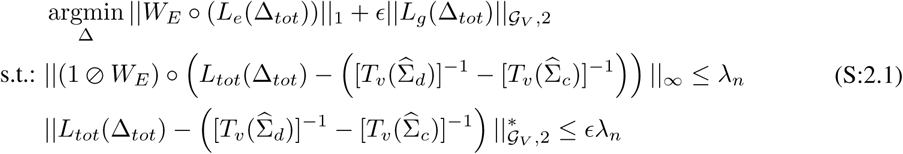

This can be rewritten as:

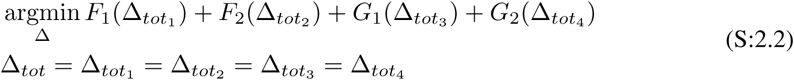

Where:

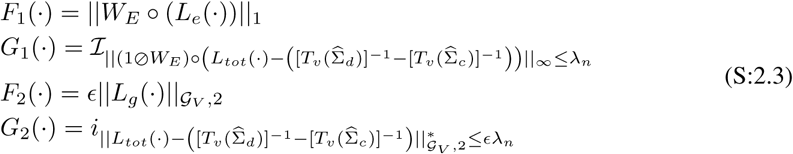

Here, *L*_*e*_, *L*_*g*_ and *L*_*tot*_ can be written as Affine Mappings. By Lemma in [],

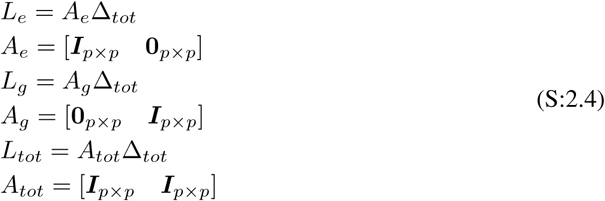

if *AA*^*T*^ = *βI*, and *h*(*x*) = *g*(*Ax*),

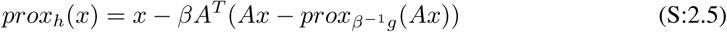

*β*_*g*_ = 1, *β*_*e*_ = 1 and *β*_*tot*_ = 2.

###### Solving for each proximal operator

**A.***F*_1_(Δ_*tot*_) = ‖*W*_*E*_ ∘ (*L*_*e*_(Δ_*tot*_))‖_1_

*L*_*e*_(Δ_*tot*_) = *A*_*e*_Δ_*tot*_ = Δ_*e*_.

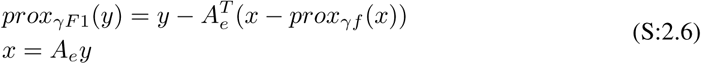

Here, 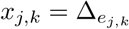.

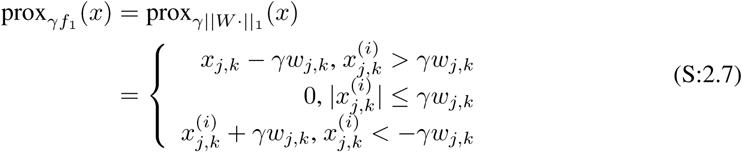

Here *j, k* = 1, …, *p*. This is an entry-wise operator (i.e., the calculation of each entry is only related to itself). This can be written in closed form:

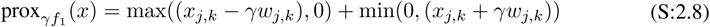

We replace this in Eq. (S:2.6).

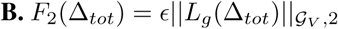 Here, *L*_*g*_(Δ_*tot*_) = *A*_*g*_Δ_*tot*_ = Δ_*g*_.

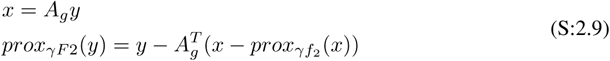

Here, 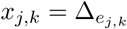.

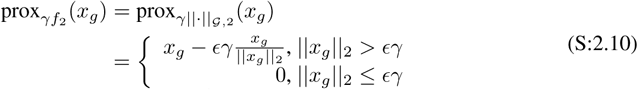

Here *g* ∈ 𝒢_𝒱_. This is a group entry-wise operator (computing a group of entries is not related to other groups). In closed form:

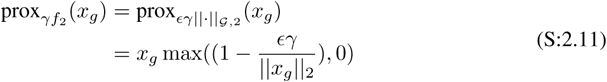

We replace this is Eq. (S:2.9).

**C.** 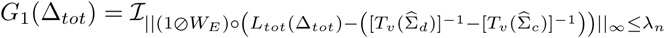 Here, *L*_*tot*_ = *A*_*tot*_Δ _*tot*_ and *A*_*tot*_ = [***I***_*p*×*p*_ ***I***_*p*×*p*_].

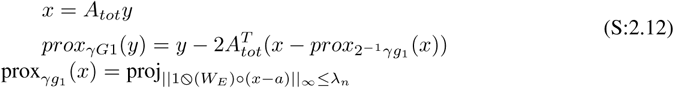

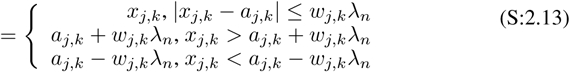

In closed form:

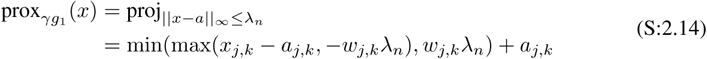

We replace this in Eq. (S:2.12).

**D.** 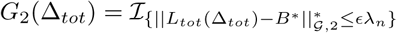 Here, *L*_*tot*_ = *A*_*tot*_Δ_*tot*_ and *A*_*tot*_ = [***I***_*p*×*p*_ ***I***_*p*×*p*_].

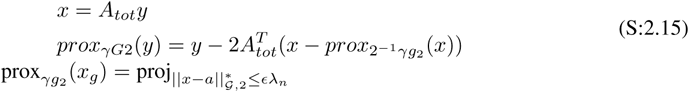

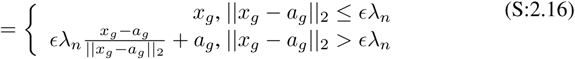

**Table S:1:** The four proximal operators

|  |  |
| --- | --- |
| $[\text{prox}_{\gamma f_1}(x)]_{j,k}^{(i)}$ | $\max((x_{j,k} - \gamma w_{j,k}), 0) + \min(0, (x_{j,k} + \gamma w_{j,k}))$ |
| $\text{prox}_{\gamma}(x_g)$ | $x_g \max((1 - \frac{\epsilon\gamma}{\ x_g\ _2}), 0)$ |
| $[\text{prox}_{\gamma f_3}(x)]_{j,k}^{(i)}$ | $\min(\max(x_{j,k} - a_{j,k}, -w_{j,k}\lambda_n), w_{j,k}\lambda_n) + a_{j,k}$ |
| $\text{prox}_{\gamma f_4}(x_g)$ | $\min(\frac{\epsilon\lambda_n}{\ x_g - a_g\ _2}, 1)(x_g - a_g) + a_g$ |

This operator is group entry-wise. In closed form:

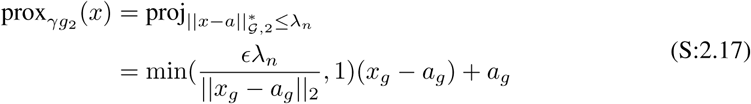

We replace this in Eq. (S:2.15).

##### S:2.2 Close form solutions if incorporating Only Edge Or Only Node Group Knowledge

In cases, where we do not have superposition structures in the differential graph estimation, we can estimate the target Δ through a closed form solution, making the method scalable to larger *p*. In detail:

###### KDiffNet-E Only Edge-level Knowledge *W*_*E*_

If additional knowledge is only available in the form of edge weights, the Eq. (S:2.1) reduces to :

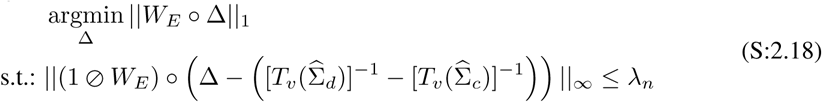

This has a closed form solution:

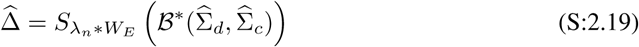

Here

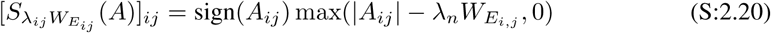

###### KDiffNet-G Only Node Groups Knowledge *G*_*V*_

If additional knowledge is only available in the form of groups of vertices 𝒢_*V*_, the Eq. (S:2.1) reduces to :

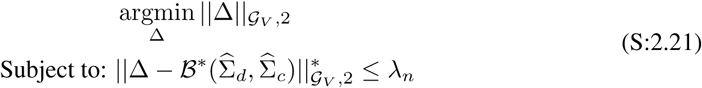

Here, we assume nodes not in any group as individual groups with cardinality= 1. The closed form solution is given by:

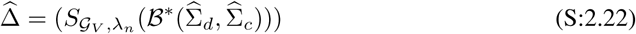

Where 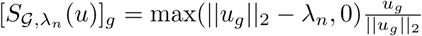 and max is the element-wise max function. Algorithm 1 shows the detailed steps of the KDiffNet estimator. Being non-iterative, the closed form solution helps KDiffNet achieve significant computational advantages.

###### Algorithm 1

KDiffNet-E and KDiffNet-G

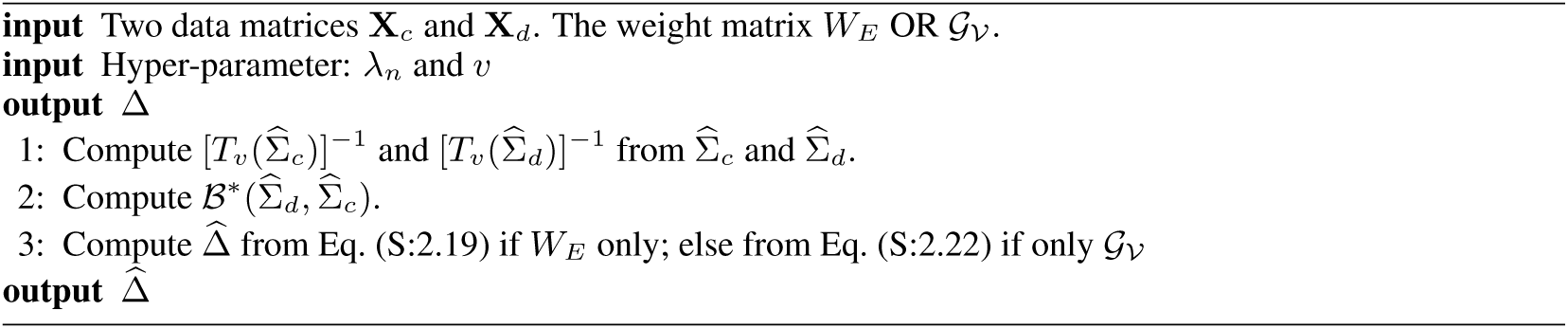

#### S:3 More Proof about kEV Norm and Its Dual Norm

##### S:3.1 Proof for kEV Norm is a norm

We reformulate kEV norm as

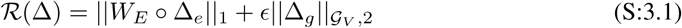

to

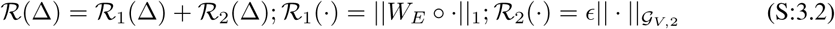

###### Theorem S:3.1.

*kEV Norm is a norm if and only if* ℛ_1_(·) *and* ℛ_2_(·) *are norms.*

*Proof.* By the following Theorem S:3.3, *R*_1_(·) is a norm. If *ϵ >* 0, *R*_2_(·) is a norm. Sum of two norms is a norm, hence kEV Norm is a norm.

###### Lemma S:3.2.

*For kEV-norm, W*_*Ej,k*_ *≠* 0 *equals to W*_*Ej,k*_ *>* 0.

*Proof.* If *W*_*Ej,k*_ *<* 0, then |*W*_*Ej,k*_Δ_*j,k*_| = | *- W*_*Ej,k*_Δ_*j,k*_|. Notice that *-W*_*Ej,k*_ *>* 0.

###### Theorem S:3.3.

ℛ_1_(·) = ||*W*_*E*_ ∘ · ||_1_ *is a norm if and only if ∀*1 *≥ j, k ≤ p, W*_*Ejk*_ *≠* 0.

*Proof. Proof.* To prove the ℛ_1_(·) = ||*W*_*E*_ ∘ · ||_1_ is a norm, by Lemma (S:4.2) we need to prove that *f* (*x*) = ||*W* ∘ *x*||_1_ is a norm function if *W*_*i,j*_ *>* 0. 1. *f* (*ax*) = ||*aW* ∘ *x*||_1_ = |*a*|||*W* ∘ *x*||_1_ = |*a*|*f* (*x*). 2. *f* (*x* + *y*) = ||*W* ∘ (*x* + *y*)||_1_ = ||*W* ∘ *x* + *W* ∘ *y*||_1_ *≤* ||*W* ∘ *x*||_1_ + ||*W* ∘ *y*||_1_ = *f* (*x*) + *f* (*y*). 3. *f* (*x*) *≥* 0. 4. If *f* (*x*) = 0, then Σ|*W*_*i,j*_*x*_*i,j*_| = 0. Since *W*_*i,j*_ *≠* 0, *x*_*i,j*_ = 0. Therefore, *x* = 0. Based on the above, *f* (*x*) is a norm function. Since summation of norm is still a norm function, ℛ_1_(·) is a norm function.

##### S:3.2 kEV Norm is a decomposable norm

We show that kEV Norm is a decomposable norm within a certain subspace, with the following structural assumptions of the true parameter Δ^*^:

###### (EV-Sparsity)

The ‘true’ parameter of Δ^*^ can be decomposed into two clear structures–{Δ_*e*_^*^and Δ_*g*_^*^}. Δ_*e*_^*^ is exactly sparse with *s* non-zero entries indexed by a support set *S*_*E*_ and Δ_*g*_^*^ is exactly sparse with 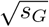 non-zero groups with atleast one entry non-zero indexed by a support set *S*_*V*_. *S*_*E*_ ⋂ *S*_*V*_ = Ø. All other elements equal to 0 (in (*S*_*E*_ ⋃ *S*_*V*_)^*c*^).

###### Definition S:3.4.

*(EV-subspace)*

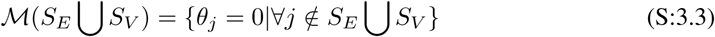

###### Theorem S:3.5.

*kEV Norm is a decomposable norm with respect to* ℳ *and* 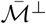

*Proof.* Assume *u* ∈ℳ and 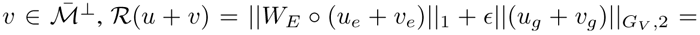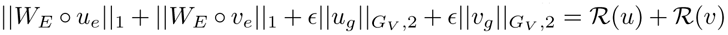. Therefore, kEV-norm is a decomposable norm with respect to the subspace pair 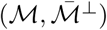.

###### Theorem S:3.6.

*Dual Norm of kEV Norm is* 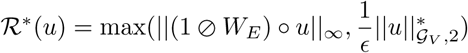.

*Proof.* Suppose 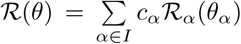, where 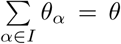. Then the dual norm ℛ^*^(·) can be derived by the following equation.

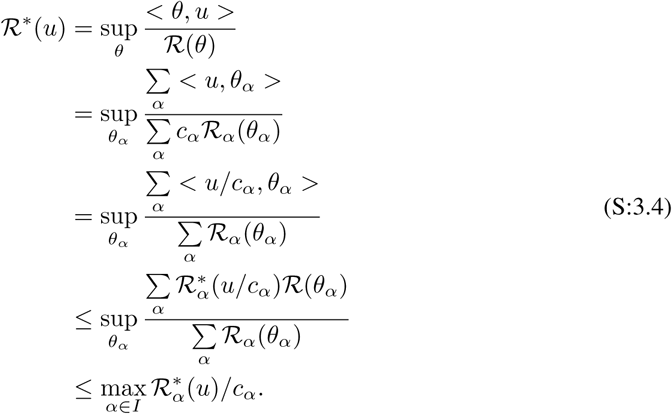

Connecting ℛ_1_(·) = ‖*W*_*E*_ · ‖_1_ and 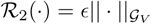. By the following Theorem S:3.7, 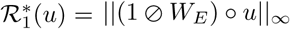. From [13], for 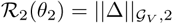, the dual norm is given by

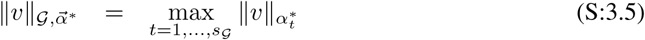

where 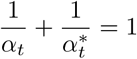 are dual exponents. where *s*_𝒢_ denotes the number of groups. As special cases of this general duality relation, this leads to a block (*∞,* 2) norm as the dual.

Hence, 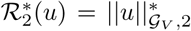. Hence, the dual norm of kEV norm is 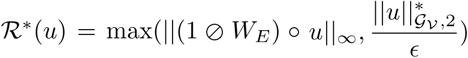.

###### Theorem S:3.7.

*The dual norm of* ‖*W*_*E*_ ∘ · ‖_1_ *is:*

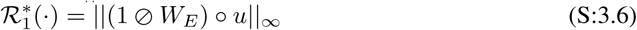

For ℛ _1_(·) = ‖*W*_*E*_ ∘‖_1_, the dual norm is given by:

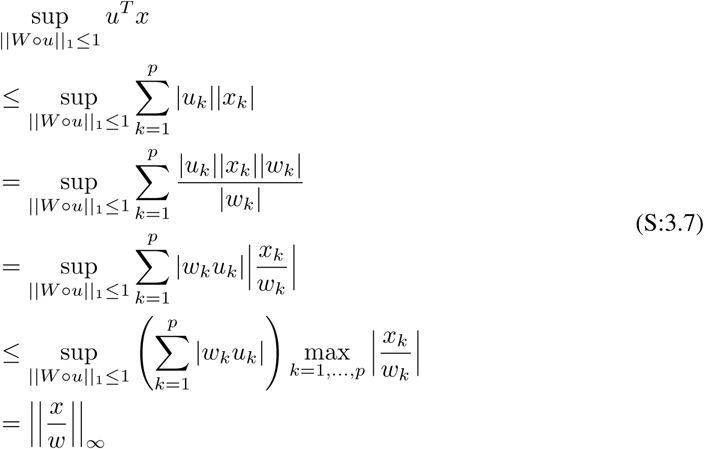

#### S:4 Appendix: More Background of Proxy Backward mapping and Theorems of *T*_*v*_ Being Invertible

Essentially the MLE solution of estimating vanilla graphical model in an exponential family distribution can be expressed as a backward mapping that computes the target model parameters from certain given moments. For instance, when learning Gaussian GM with vanilla MLE, the backward mapping is 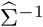 that estimates Ω from the sample covariance matrix (moment) 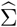. However, this backward mapping is normally not well-defined in high-dimensional settings. In the case of GGM, when given the sample covariance 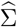, we cannot just compute the vanilla MLE solution as 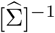 when high-dimensional since 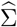 is rank-deficient when *p > n*. Therefore Yang et al. [24] proposed to use carefully constructed proxy backward maps for Eq. (2.4) that are both available in closed-form, and well-defined in high-dimensional settings for exponential GM models. For instance, 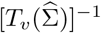 in Eq. (2.3) is the proxy backward mapping [24] used for GGM.

##### S:4.1 Backward mapping for an exponential-family distribution

The solution of vanilla graphical model MLE can be expressed as a backward mapping[21] for an exponential family distribution. It estimates the model parameters (canonical parameter *θ*) from certain (sample) moments. We provide detailed explanations about backward mapping of exponential families, backward mapping for Gaussian special case and backward mapping for differential network of GGM in this section.

###### Backward mapping

Essentially the vanilla graphical model MLE can be expressed as a backward mapping that computes the model parameters corresponding to some given moments in an exponential family distribution. For instance, in the case of learning GGM with vanilla MLE, the backward mapping is 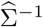 that estimates Ω from the sample covariance (moment) 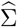.

Suppose a random variable *X* ∈ ℝ^*p*^ follows the exponential family distribution:

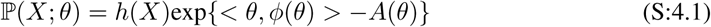

Where *θ* ∈ Θ *⊂*ℝ^*d*^ is the canonical parameter to be estimated and Θ denotes the parameter space. *ϕ*(*X*) denotes the sufficient statistics as a feature mapping function *ϕ* : ℝ^*p*^ *ν*ℝ^*d*^, and *A*(*θ*) is the log-partition function. We then define mean parameters *v* as the expectation of *ϕ*(*X*): *v*(*θ*) := 𝔼[*ϕ*(*X*)], which can be the first and second moments of the sufficient statistics *ϕ*(*X*) under the exponential family distribution. The set of all possible moments by the moment polytope:

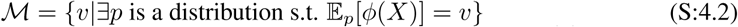

Mostly, the graphical model inference involves the task of computing moments *v*(*θ*) ∈ℳ given the canonical parameters 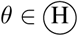. We denote this computing as **forward mapping** :

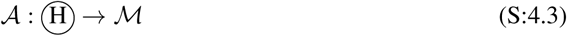

The learning/estimation of graphical models involves the task of the reverse computing of the forward mapping, the so-called **backward mapping** [21]. We denote the interior of ℳ as ℳ^0^. **backward mapping** is defined as:

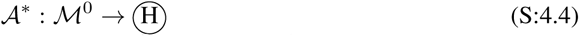

which does not need to be unique. For the exponential family distribution,

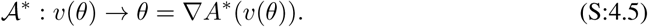

Where 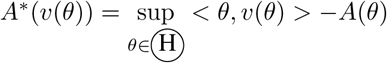.

###### Backward Mapping: Gaussian Case

If a random variable *X* ∈ ℝ^*p*^ follows the Gaussian Distribution *N* (*µ,* Σ). then 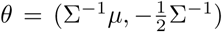. The sufficient statistics 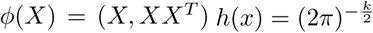, and the log-partition function

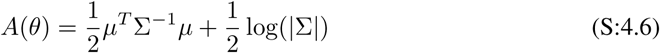

When performing the inference of Gaussian Graphical Models, it is easy to estimate the mean vector *v*(*θ*), since it equals to 𝔼[*X, XX*^*T*^].

When learning the GGM, we estimate its canonical parameter *θ* through vanilla MLE. Because Σ^−1^ is one entry of *θ* we can use the backward mapping to estimate Σ^−1^.

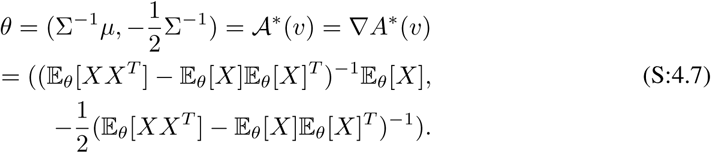

By plugging in Eq. (S:4.6) into Eq. (S:4.5), we get the backward mapping of Ω as 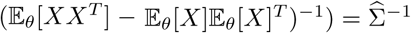, easily computable from the sample covariance matrix.

##### S:4.2 Backward Mapping for Differential GGM

When the random variables *X*_*c*_, *X*_*d*_ ∈ℝ^*p*^ follows the Gaussian Distribution *N* (*µ*_*c*_, Σ_*c*_) and *N* (*µ*_*d*_, Σ_*d*_), their density ratio (defined by [12]) essentially is a distribution in exponential families:

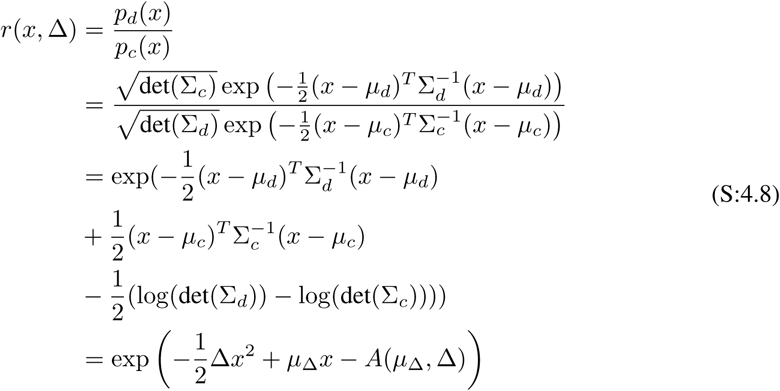

Here 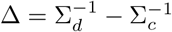 and 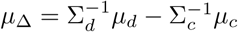.

The log-partition function

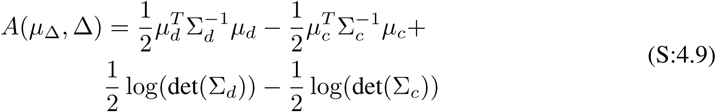

The canonical parameter

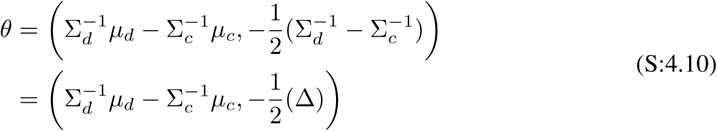

The sufficient statistics *ϕ*([*X*_*c*_, *X*_*d*_]) and the log-partition function *A*(*θ*):

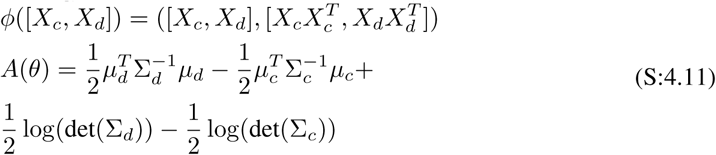

And *h*(*x*) = 1.

Now we can estimate this exponential distribution (*θ*) through vanilla MLE. By plugging Eq. (S:4.11) into Eq. (S:4.5), we get the following backward mapping via the conjugate of the log-partition function:

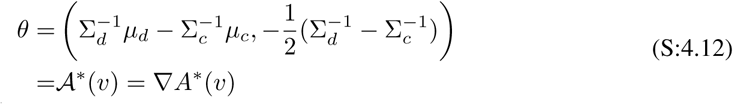

The mean parameter vector *v*(*θ*) includes the moments of the sufficient statistics *ϕ*() under the exponential distribution. It can be easily estimated through 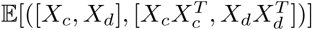.

Therefore the backward mapping of *θ* becomes,

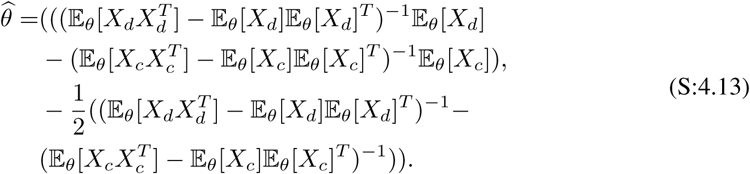

Because the second entry of the canonical parameter *θ* is 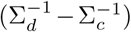, we get the backward mapping of Δ as

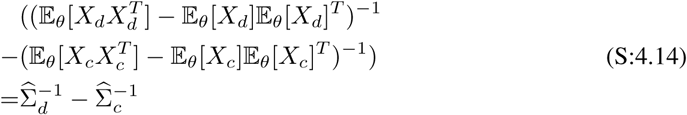

This can be easily inferred from two sample covariance matrices 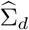 and 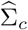 (Att: when under low-dimensional settings).

##### S:4.3 Theorems of Proxy Backward Mapping *T*_*v*_ Being Invertible

Based on [24] for any matrix A, the element wise operator *T*_*v*_ is defined as:

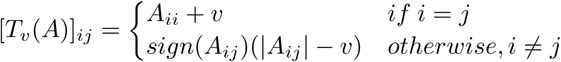

Suppose we apply this operator *T*_*v*_ to the sample covariance matrix 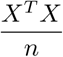 to obtain 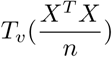. Then, 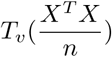 under high dimensional settings will be invertible with high probability, under the following conditions:

**Condition-1** (Σ-Gaussian ensemble) Each row of the design matrix *X* ∈ℝ^*n*×*p*^ is i.i.id sampled from *N* (0, Σ).

**Condition-2** The covariance Σ of the Σ-Gaussian ensemble is strictly diagonally dominant: for all row i, *δ*_*i*_ := Σ_*ii*_ *-* Σ_*j≠i*_ *≥ δ*_*min*_ *>* 0 where *δ*_*min*_ is a large enough constant so that 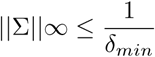.

This assumption guarantees that the matrix 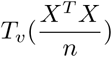 is invertible, and its induced 𝓁_*∞*_ norm is well bounded. Then the following theorem holds:

###### Theorem S:4.1.

*Suppose Condition-1 and Condition-2 hold. Then for any v ≥* 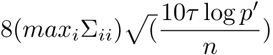, *the matrix* 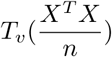 *is invertible with probability at least* 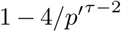 *for p′* := *max*{*n, p*} *and any constant τ >* 2.

##### S:4.4 Useful lemma(s) of Error Bounds of Proxy Backward Mapping *T*_*v*_

###### Lemma S:4.2.

*(Theorem 1 of [16]). Let δ be* 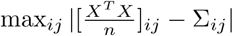. *Suppose that ν >* 2*δ. Then, under the conditions (C-Sparse*Σ*), and as ρ*_*v*_() *is a soft-threshold function, we can deterministically guarantee that the spectral norm of error is bounded as follows:*

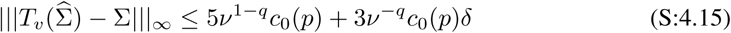

###### Lemma S:4.3.

*(Lemma 1 of [15]). Let* 𝒜 *be the event that*

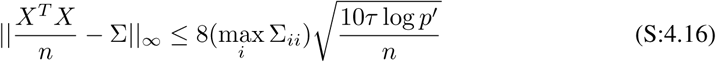

*where p′* := max(*n, p*) *and τ is any constant greater than 2. Suppose that the design matrix X is i.i.d. sampled from* Σ*-Gaussian ensemble with n ≥* 40 max_*i*_ Σ_*ii*_. *Then, the probability of event* 𝒜 *occurring is at least* 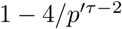.

#### S:5 Theoretical Analysis of Error Bounds

##### S:5.1 Background: Error bounds of Elementary Estimators

KDiffNet formulations are special cases of the following generic formulation for the elementary estimator.

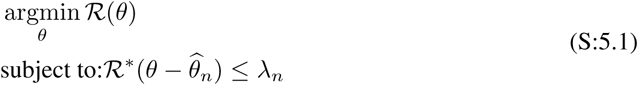

Where ℛ^*^(·) is the dual norm of ℛ(·),

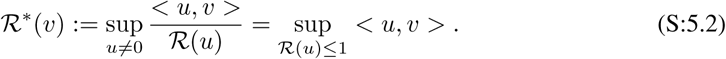

Following the unified framework [13], we first decompose the parameter space into a subspace pair 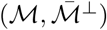, where 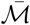 is the closure of *ℳ*. Here 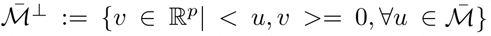. *ℳ* is the **model subspace** that typically has a much lower dimension than the original high-dimensional space. 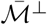 is the **perturbation subspace** of parameters. For further proofs, we assume the regularization function in Eq. (S:5.1) is **decomposable** w.r.t the subspace pair 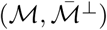.

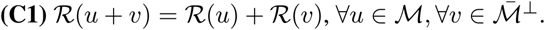

[13] showed that most regularization norms are decomposable corresponding to a certain subspace pair.

###### Definition S:5.1.

*Subspace Compatibility Constant*

*Subspace compatibility constant is defined as* 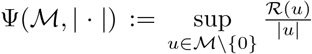 *which captures the relative value between the error norm* | · | *and the regularization function* ℛ(·).

For simplicity, we assume there exists a true parameter *θ*^*^ which has the exact structure w.r.t a certain subspace pair. Concretely:

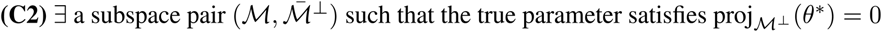

Then we have the following theorem.

###### Theorem S:5.2.

*Suppose the regularization function in Eq. (S:5.1) satisfies condition* ***(C1)***, *the true parameter of Eq. (S:5.1) satisfies condition* ***(C2)***, *and λ*_*n*_ *satisfies that* 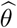. *Then, the optimal solution* 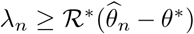 *of Eq. (S:5.1) satisfies:*

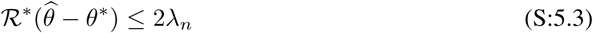

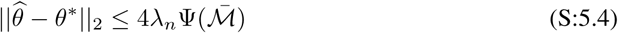

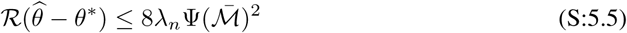

*Proof.* Let 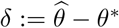 be the error vector that we are interested in.

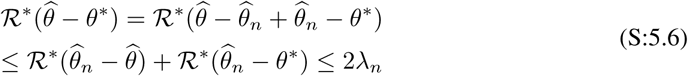

By the fact that 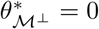, and the decomposability of *ℛ* with respect to 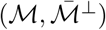

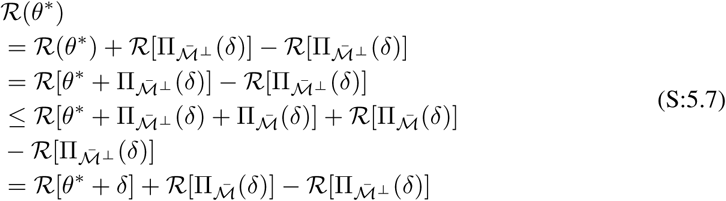

Here, the inequality holds by the triangle inequality of norm. Since Eq. (S:5.1) minimizes 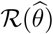, we have 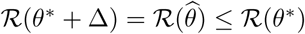. Combining this inequality with Eq. (S:5.7), we have:

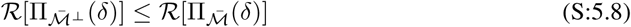

Moreover, by Hölder’s inequality and the decomposability of *ℛ*;(·), we have:

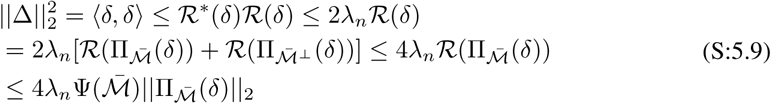

where 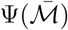 is a simple notation for 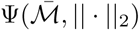.

Since the projection operator is defined in terms of ‖· ‖_2_ norm, it is non-expansive: 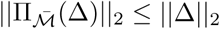. Therefore, by Eq. (S:5.9), we have:

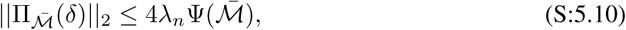

and plugging it back to Eq. (S:5.9) yields the error bound Eq. (S:5.4).

Finally, Eq. (S:5.5) is straightforward from Eq. (S:5.8) and Eq. (S:5.10).

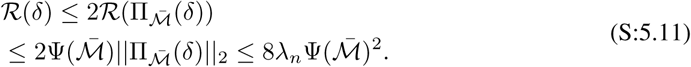

##### S:5.2 Error Bounds of KDiffNet

Theorem S:5.2, provides the error bounds via *λ*_*n*_ with respect to three different metrics. In the following, we focus on one of the metrics, Frobenius Norm to evaluate the convergence rate of our KDiffNet estimator.

###### Theorem S:5.3.

*Assuming the true parameter* Δ^*^ *satisfies the conditions* ***(C1)(C2)*** *and* 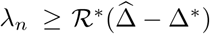, *then the optimal point* 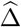 *has the following error bounds:*

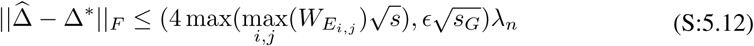

Proof: KDiffNet uses ℛ(·)= ‖*W*_*E*_°·‖_1_ +*ϵ* ‖·‖_𝒢,2_ because it is a superposition of two norms: ℛ_1_‖ *W*_*E*_°·‖_1_ and ℛ_2_ = *ϵ*‖·‖_𝒢,2_ Based on the results in[13], 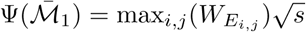 and 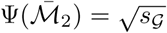, where *s* is the number of nonzero entries in Δ and *s*_*G*_ is the number of groups in which there exists at least one nonzero entry. Therefore, 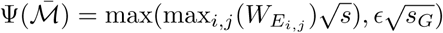 Hence, Using this in Equation Eq. (S:5.4), 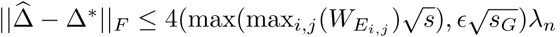.

###### S:5.2.2 Proof of Corollary (2.1)-Derivation of the KDiffNet error bounds

To derive the convergence rate for KDiffNet, we introduce the following two sufficient conditions on the ∑_*c*_ and ∑_*d*_, to show that the proxy backward mapping 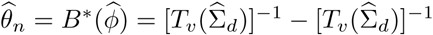 is well-defined[23]:

**(C-MinInf−∑)** The true 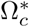 and 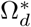 of Eq. (2.1) have bounded induced operator norm, i.e., 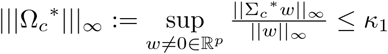 and 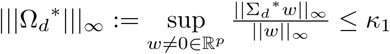.

**(C-Sparse-∑):** The two true covariance matrices 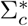 and 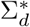 are “approximately sparse” (following [2]). For some constant 0 ≤ *q <* 1 and 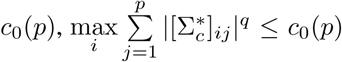 and 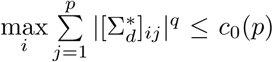.^2^

We additionally require 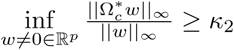 and 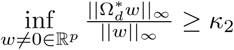.

We assume the true parameters 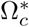 and 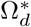 satisfies **C-MinInf**∑ and **C-Sparse**∑ conditions.

Using the above theorem and conditions, we have the following corollary for convergence rate of KDiffNet (Att: the following corollary is the same as the Corollary 2.1 in the main draft. We repeat it here to help readers read the manuscript more easily):

###### Corollary S:5.4.

*In the high-dimensional setting, i.e.*, 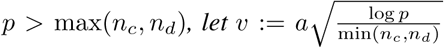. *Then for* 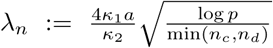 *and* min(*n*_*c*_, *n*_*d*_) *> c* log *p, with a probability of at least* 1 −2*C*_1_ exp(*−C*_2_*p* log(*p*)), *the estimated optimal solution* 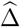 *has the following error bound:*

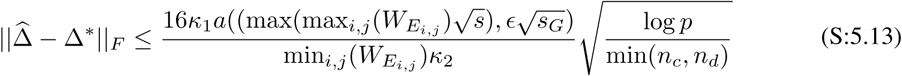

*where a, c, κ*_1_ *and κ*_2_ *are constants.*

*Proof.* In the following proof, we first prove 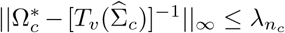. Here 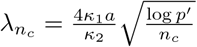 and *p*^′^ = max(*p, n*_*c*_)

The condition (C-Sparse∑) and condition (C-MinInf∑) also hold for 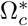 and 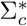. In order to utilize Theorem (S:5.3) for this specific case, we only need to show that 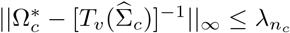 for the setting of 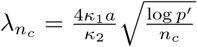:

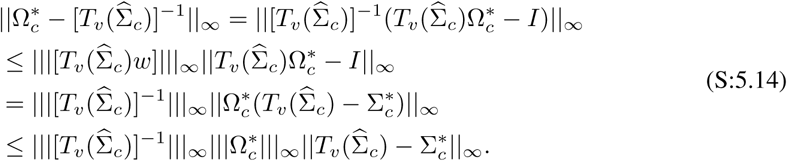

We first compute the upper bound of 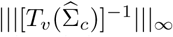. By the selection *v* in the statement, Lemma (S:4.2) and Lemma (S:4.3) hold with probability at least 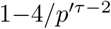. Armed with Eq. (S:4.15), we use the triangle inequality of norm and the condition (C-Sparse∑): for any *w*,

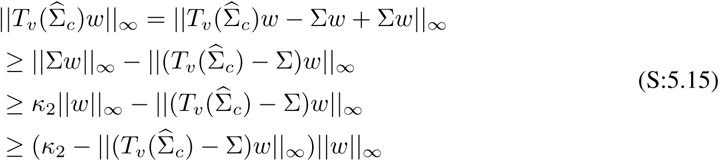

Where the second inequality uses the condition (C-Sparse∑). Now, by Lemma (S:4.2) with the selection of *v*, we have

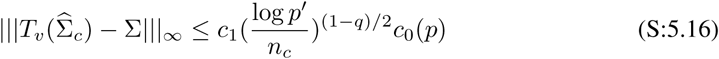

where *c*_1_ is a constant related only on *τ* and max_*i*_ ∑_*ii*_. Specifically, it is defined as 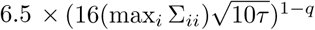. Hence, as long as 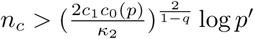 as stated, so that 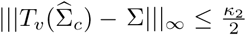, we can conclude that 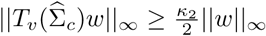, which implies 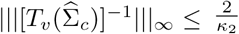.

The remaining term in Eq. (S:5.14) is 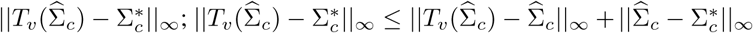. By construction of *T*_*v*_(·) in (C-Thresh) and by Lemma (S:4.3), we can confirm that 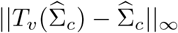 as well as 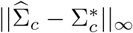 can be upper-bounded by *v*.

Similarly, the 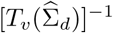 has the same result.

Finally,

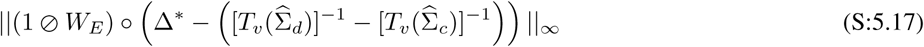

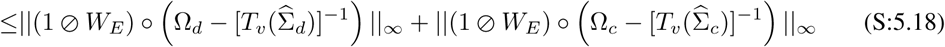

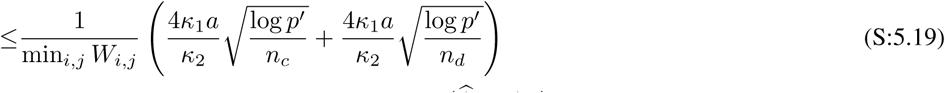

Because by Theorem S:5.3, we know if 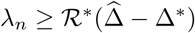,

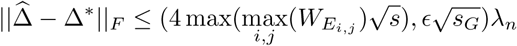

Suppose *p* > max(*n*_*c*_, *n*_*d*_) we have that

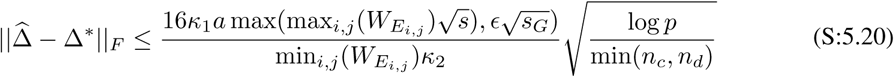

By combining all together, we can confirm that the selection of *λ*_*n*_ satisfies the requirement of Theorem (S:5.3), which completes the proof.

#### S:6 More about Experiments

##### S:6.1 Experimental Setup

*The hyper-parameters in our experiments are v, λ*_*n*_, *ϵ* and *λ*_2_. *In detail:*

- To compute the proxy backward mapping in (S:2.1), DIFFEE, and JEEK we vary *v* for soft-thresholding *v* from the set {0.001*i*|*i* = 1, 2, *…,* 1000} (to make *T*_*v*_(∑_*c*_) and *T*_*v*_(∑_*d*_) invertible).
- *λ*_*n*_ is the hyper-parameter in our KDiffNet formulation. According to our convergence rate analysis in Section 2.6, 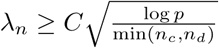, we choose *λ*_*n*_ from a range of 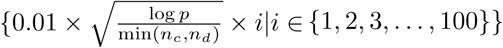. For KDiffNet-G case, we tune over *λ*_*n*_ from a range of 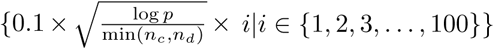. We use the same range to tune *λ*_1_ for SDRE. Tuning for NAK is done by the package itself.
- *ϵ*: For KDiffNet-EG experiments, we tune *ϵ* ∈{0.0001, 0.01, 1, 100}.
- *λ*_2_ controls individual graph’s sparsity in JGLFUSED. We choose *λ*_1_ = 0.0001 (a very small value) for all experiments to ensure only the differential network is sparse.

###### Evaluation Metrics

- F1-score: We use the edge-level F1-score as a measure of the performance of each method. 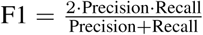, where 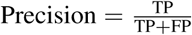 and 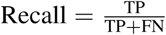. The better method achieves a higher F1-score. We choose the best performing *λ*_*n*_ using validation and report the performance on a test dataset.
- Time Cost: We use the execution time (measured in seconds or log(seconds)) for a method as a measure of its scalability. The better method uses less time^3^

##### S:6.2 Simulation Dataset Generation

We first use simulation to evaluate KDiffNet for improving differential structure estimation by making use of extra knowledge. We generate simulated datasets with a clear underlying differential structure between two conditions, using the following method:

###### Data Generation for Edge Knowledge (KE)

Given a known weight matrix *W*_*E*_ (e.g., spatial distance matrix between *p* brain regions), we set *W*^*d*^ = *inv.logit*(−*W*_*E*_). We use the assumption that higher the value of *W*_*ij*_, lower the probability of that edge to occur in the true precision matrix. This is motivated by the role of spatial distance in brain connectivity networks: farther regions are less likely to be connected and vice-versa. We select different levels in the matrix *W*^*d*^, denoted by *s*, where if 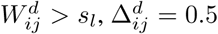, else 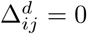, where Δ^*d*^ ∈ ∝^*p×p*^. We denote by *s* as the sparsity, i.e. the number of non-zero entries in Δ^*d*^· *B*_*I*_ is a random graph with each edge 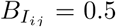 with probability *p*. *δ*_*c*_ and *δ*_*d*_ are selected large enough to guarantee positive definiteness.

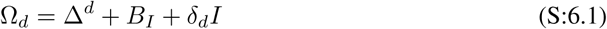

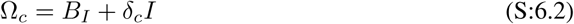

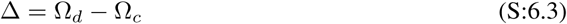

There is a clear differential structure in Δ = Ω_*d*_ − Ω_*c*_, controlled by Δ^*d*^. To generate data from two conditions that follows the above differential structure, we generate two blocks of data samples following Gaussian distribution using 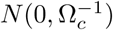 and 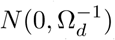. We only use these data samples to approximate the differential GGM to compare to the ground truth Δ.

###### Data Generation for Vertex Knowledge (KG)

In this case, we simulate the case of extra knowledge of nodes in known groups. Let the node group size, i.e., the number of nodes with a similar interaction pattern in the differential graph be *m*. We select the block diagonals of size *m* as groups in Δ^*g*^. If two variables *i, j* are in a group *g*^′^, in 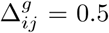, else 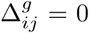, where Δ^*g*^ ∈ ℝ^*p×p*^. We denote by *s*_*G*_ as the number of groups in Δ^*g*^. *B*_*I*_ is a random graph with each edge 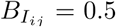 with probability *p*.

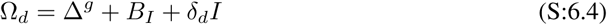

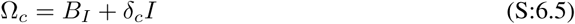

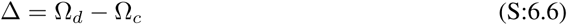

*δ*_*c*_ and *δ*_*d*_ are selected large enough to guarantee positive definiteness. We generate two blocks of data samples following Gaussian distribution using 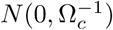 and 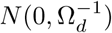.

###### Data Generation for both Edge and Vertex Knowledge (KEG)

In this case, we simulate the case of overlapping group and edge knowledge. Let the node group size,i.e., the number of nodes with a similar interaction pattern in the differential graph be *m*. We select the block diagonals of size *m* as groups in Δ^*g*^. If two variables *i, j* are in a group *g*^′^, in 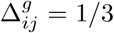, else 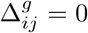, where Δ^*g*^ ∈ ℝ^*p×p*^.

For the edge-level knowledge component, given a known weight matrix *W*_*E*_, we set *W*^*d*^ = *inv.logit*(*-W*_*E*_). Higher the value of 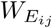, lower the value of 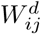, hence lower the probability of that edge to occur in the true precision matrix. We select different levels in the matrix *W*^*d*^, denoted by *s*, where if 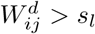, we set 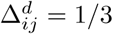, else 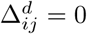. We denote by *s* as the number of non-zero entries in Δ^*d*^. *B*_*I*_ is a random graph with each edge 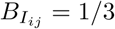 with probability *p*.

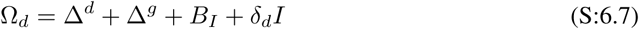

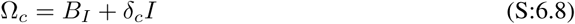

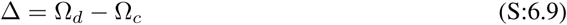

*δ*_*c*_ and *δ*_*d*_ are selected large enough to guarantee positive definiteness. Similar to the previous case, we generate two blocks of data samples following Gaussian distribution using 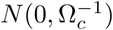 and 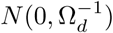. We only use these data samples to approximate the differential GGM to compare to the ground truth Δ.

##### S:6.3 Simulation Experiment Results

We consider three different types of known edge knowledge *W*_*E*_ generated from the spatial distance between different brain regions and simulate groups to represent related anatomical regions. These three are distinguished by different *p* = 116, 160, 246 representing spatially related brain regions. We generate three types of datasets:Data-EG (having both edge and vertex knowledge), Data-G(with edge-level extra knowledge) and Data-V(with known node groups knowledge). We generate two blocks of data samples ***X***_*c*_ and ***X***_*d*_ following Gaussian distribution using 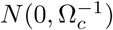 and 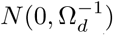. We use these data samples to estimate the differential GGM to compare to the ground truth Δ. The details of the simulation are in Section S:6.2. We vary the sparsity of the true differential graph (*s*) and the number of control and case samples (*n*_*c*_ and *n*_*d*_ respectively) used to estimate the differential graph. For each case of *p*, we vary *n*_*c*_ and *n*_*d*_ in {*p/*2, *p/*4, *p,* 2*p*} to account for both high dimensional and low dimensional cases. The sparsity of the underlying differential graph is controlled by *s* = {0.125, 0.25, 0.375, 0.5} and *s*_*G*_ as explained in Section S:6.2. This results in 126 different datasets representing diverse settings: different number of dimensions *p*, number of samples *n*_*c*_ and *n*_*d*_, multiple levels of sparsity *s* and number of groups *s*_*G*_ of the differential graph for both KE and KEG data settings.

###### Edge and Vertex Knowledge (KEG)

We use KDiffNet (Algorithm 1) to infer the differential structure in this case.

Figure S:2(a) shows the performance in terms of F1 Score of KDiffNet in comparison to the baselines for *p* = 116, corresponding to 116 regions of the brain. KDiffNet outperforms the best baseline in each case by an average improvement of 414%. KDiffNet-EG does better than JEEK and NAK that can model the edge information but cannot include group information. SDRE and DIFFEE are direct estimators but perofrm poorly indicating that adding additional knowledge aids differential network estimation. JGLFUSED performs the worst on all cases. We list the detailed results in Section S:6.5.

**Figure S:2:**
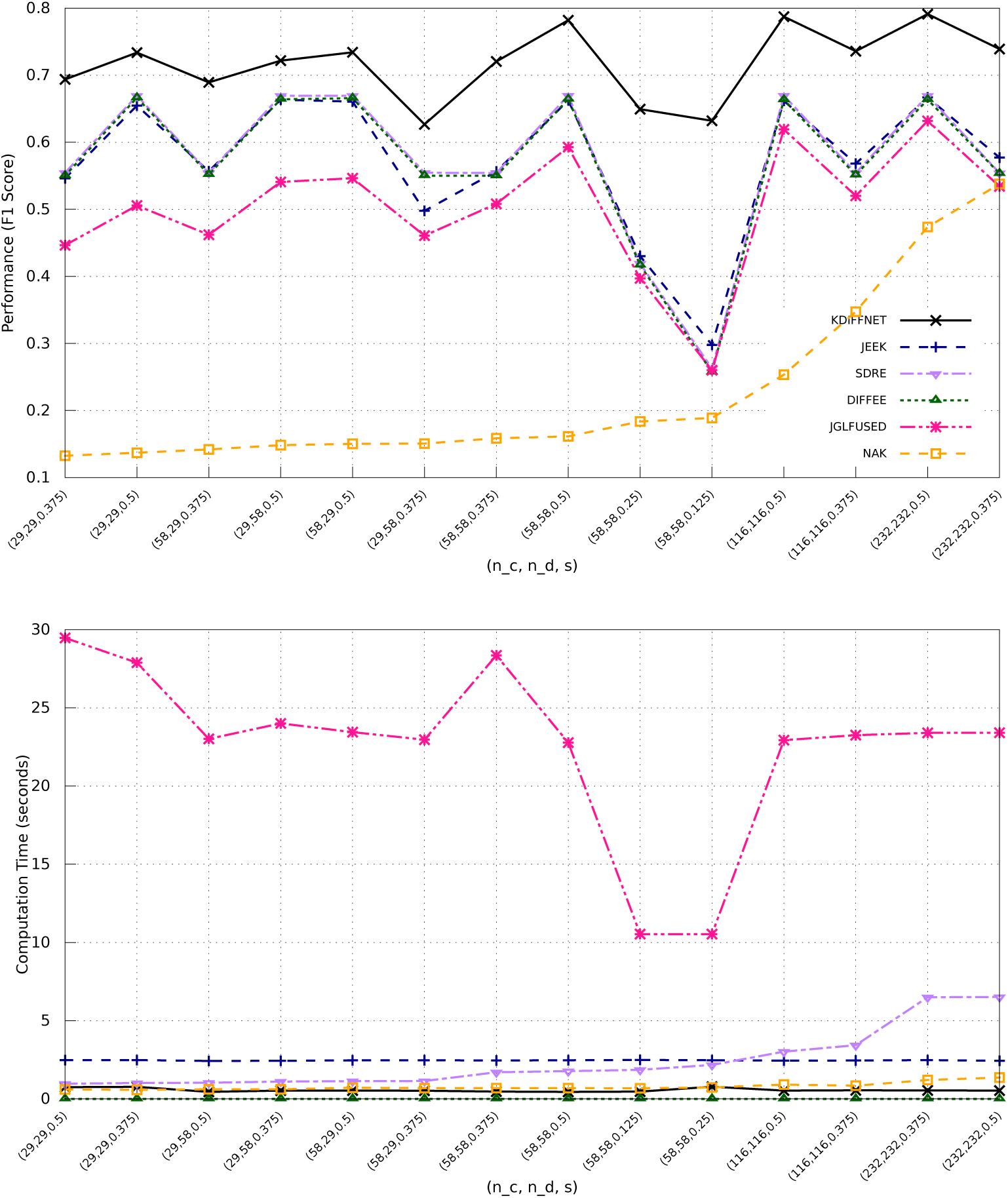
KDiffNet Edge and Vertex Knowledge Simulation Results for *p* = 116 for different settings of *n*_*c*_, *n*_*d*_ and *s*: (a) The test F1-score and (b) The average computation time (measured in seconds) per *λ*_*n*_ for KDiffNet and baseline methods.

Figure S:2(b) shows the average computation cost per *λ*_*n*_ of each method measured in seconds. In all settings, KDiffNet has lower computation cost than JEEK, SDRE and JGLFUSED in different cases of varying *n*_*c*_ and *n*_*d*_, as well as with different sparsity of the differential network. KDiffNet is on average 24× faster than the best performing baseline. It is slower than DIFFEE owing to DIFFEE’s non-iterative closed form solution, however, DIFFEE does not have good prediction performance. Note that *B*^*^() in KDiffNet, JEEK and DIFFEE and the kernel term in SDRE are precomputed only once prior to tuning across multiple *λ*_*n*_. In Figure S:3(a), we plot the test F1-score for simulated datasets generated using *W* with *p* = 160, representing spatial distances between different 160 regions of the brain. This represents a larger and different set of spatial brain regions. In *p* = 160 case, KDiffNet outperforms the best baseline in each case by an average improvement of 928%. Including available additional knowledge is clearly useful as JEEK does relatively better than the other baselines. JGLFUSED performs the worst on all cases. Figure S:3(b) shows the computation cost of each method measured in seconds for each case. KDiffNet is on average 37× faster than the best performing baseline.

**Figure S:3:**
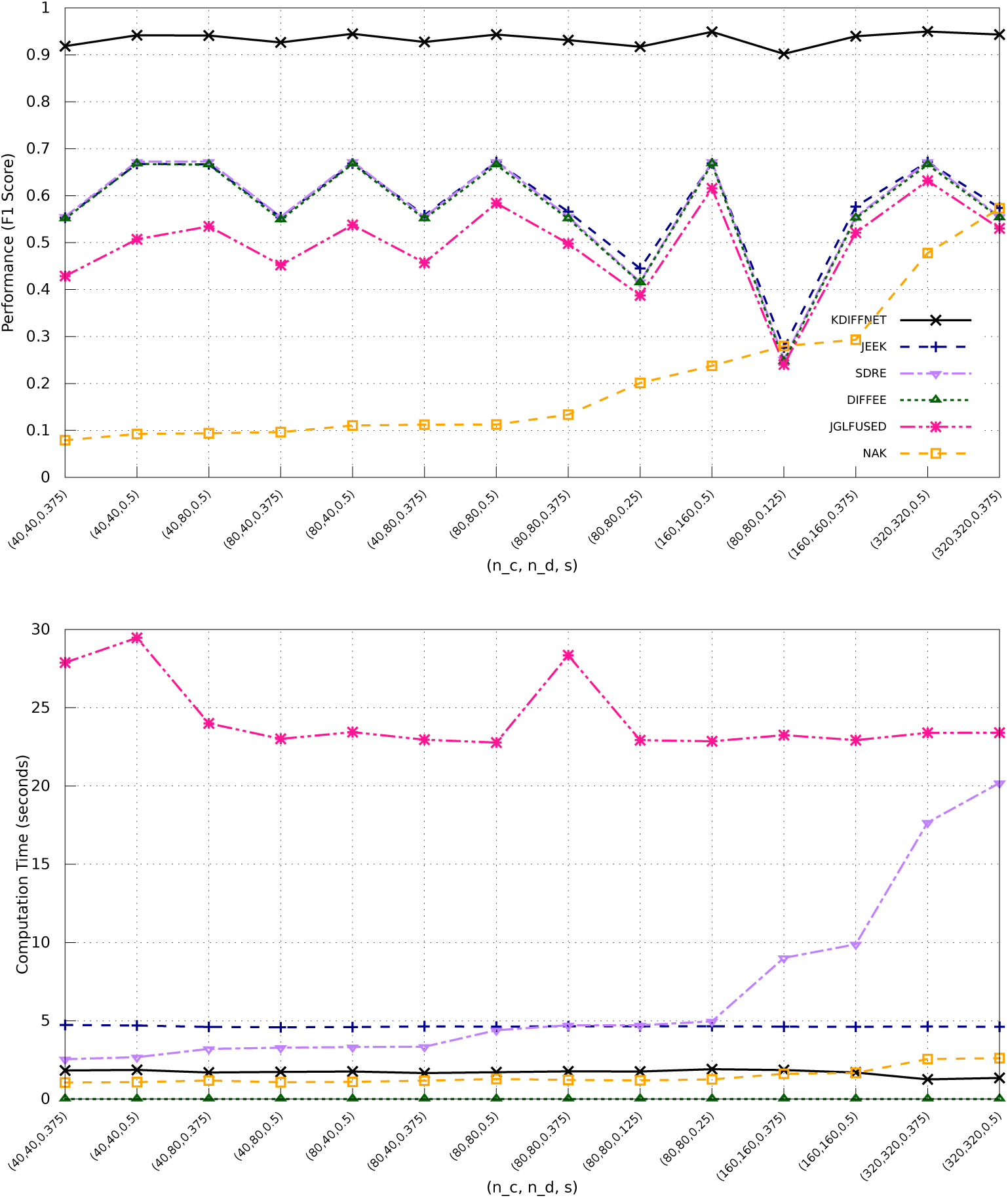
KDiffNet Edge and Vertex Knowledge Simulation Results for *p* = 160 for different settings of *n*_*c*_, *n*_*d*_ and *s*: (a) The test F1-score and (b) The average computation time (measured in seconds) per *λ*_*n*_ for KDiffNet and baseline methods.

**Figure S:4:**
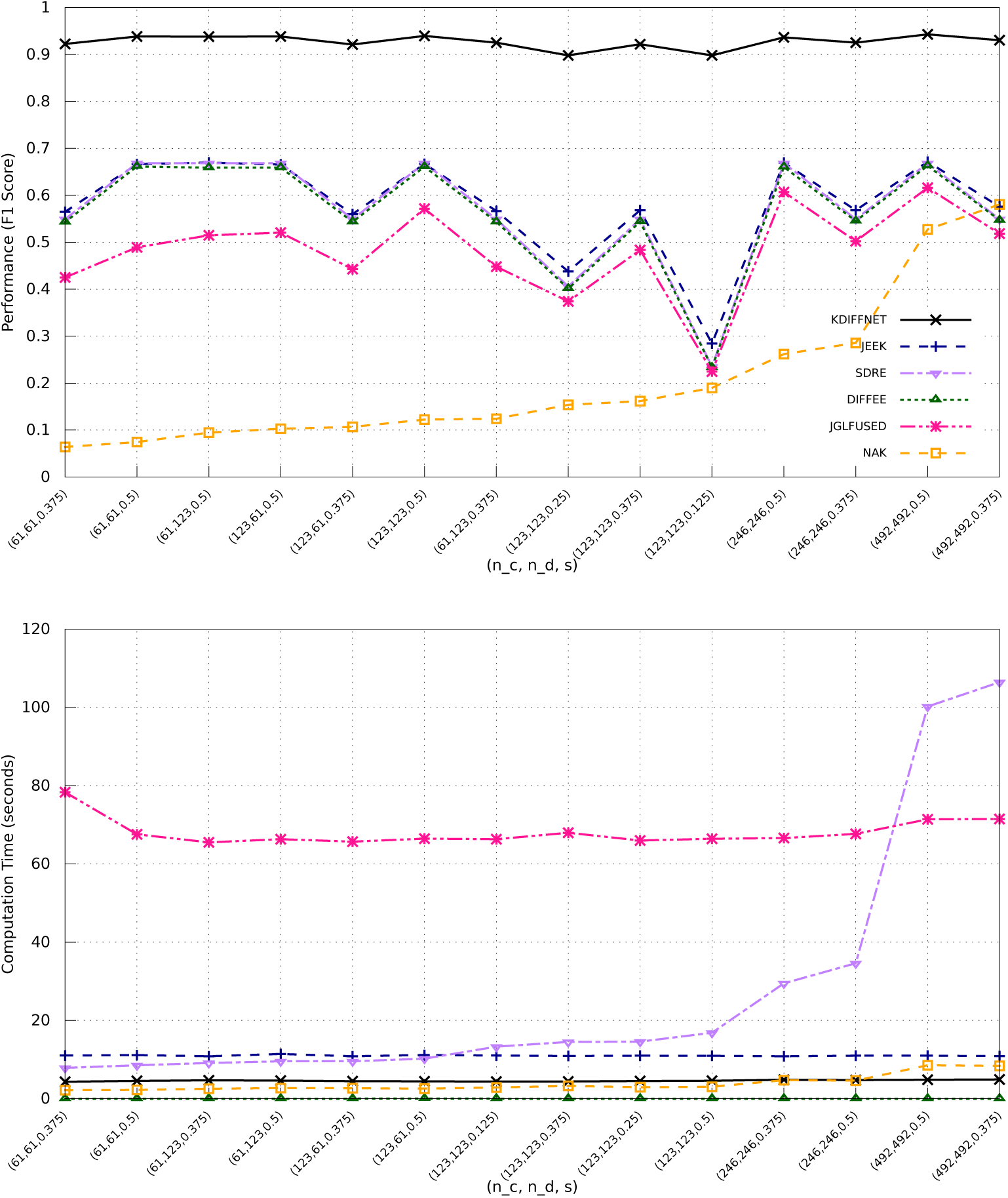
KDiffNet Edge and Vertex Knowledge Simulation Results for *p* = 246 for different settings of *n*_*c*_, *n*_*d*_ and *s*: (a) The test F1-score and (b) The average computation time (measured in seconds) per *λ*_*n*_ for KDiffNet and baseline methods

In Figure S:4(a), we plot the test F1-score for simulated datasets generated using a larger *W*_*E*_ with *p* = 246, representing spatial distances between different 246 regions of the brain. This represents a larger and different set of spatial brain regions. In this case, KDiffNet outperforms the best baseline in each case by an average improvement of 1400% relative to the best performing baseline. In this case as well, including available additional knowledge is clearly useful as JEEK does relatively better than the other baselines, which do not incorporate available additional knowledge. JGLFUSED again performs the worst on all cases. Figure S:4(b) shows the computation cost of each method measured in seconds for each case. In all cases, KDiffNet has the least computation cost in different settings of the data generation. KDiffNet is on average 20× faster than the best performing baseline. For detailed results, see Section S:6.5.

We cannot compare Diff-CLIME as it takes more than 2 days to finish *p* = 246 case.

###### Edge Knowledge (KE)

Given known *W*_*E*_, we use KDiffNet-E to infer the differential structure in this case.

**Figure S:5:**
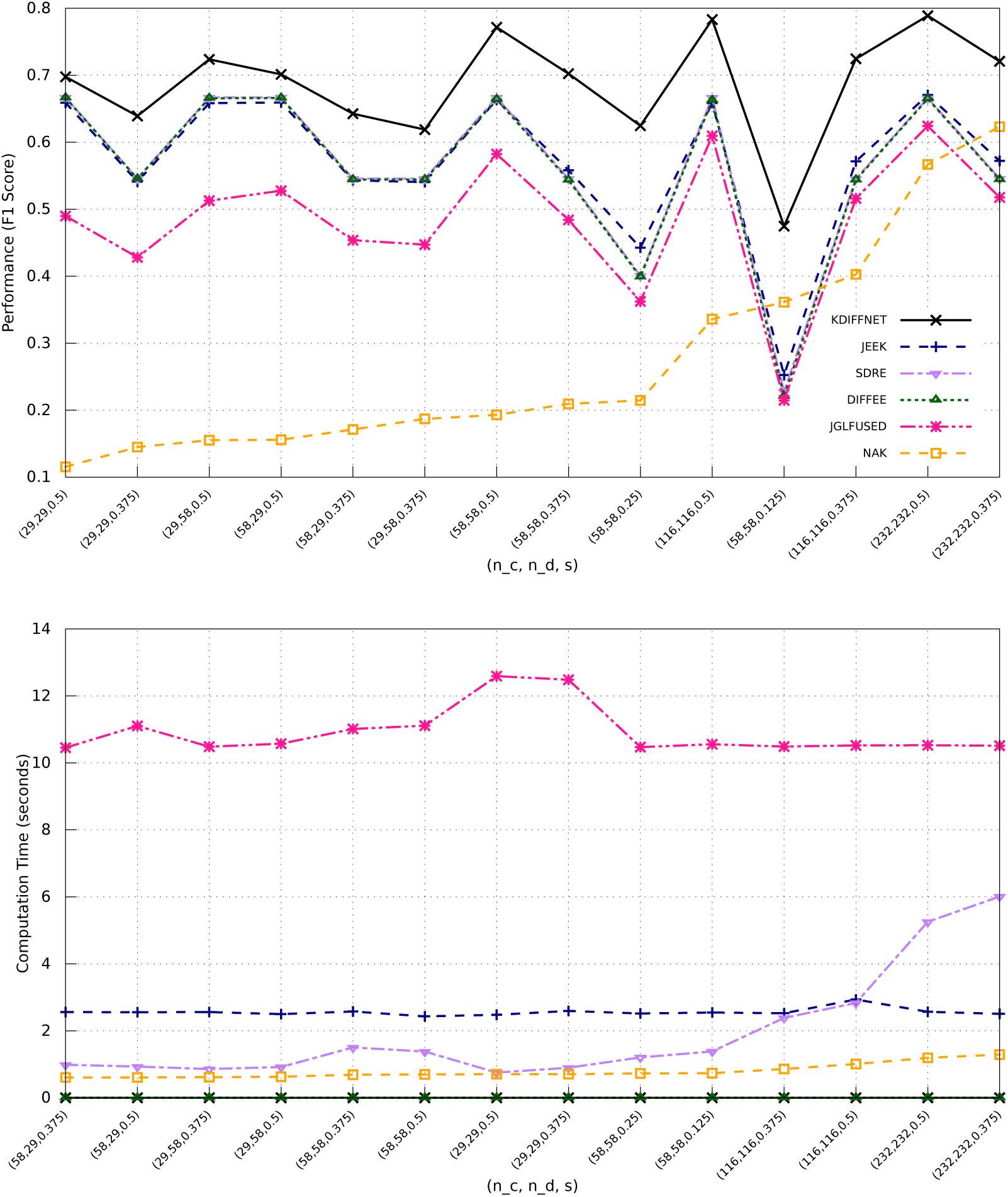
(a) shows the performance in terms of F1-Score of KDiffNet-E in comparison to the baselines for *p* = 116, corresponding to 116 spatial regions of the brain. In *p* = 116 case, KDiffNet-E outperforms the best baseline in each case by an average improvement of 23%. While JEEK, DIFFEE and SDRE perform similar to each other, JGLFUSED performs the worst on all cases.

Figure S:5: KDiffNet-E Simulation Results for *p* = 116 for different settings of *n*_*c*_, *n*_*d*_ and *s*: (a) The test F1-score and (b) The average computation time (measured in seconds) per *λ*_*n*_ for KDiffNet-E and baseline methods.

Figure S:5(b) shows the computation cost of each method measured in seconds for each case. In all cases, KDiffNet-E has the least computation cost in different cases of varying *n*_*c*_ and *n*_*d*_, as well as with different sparsity of the differential network. For *p* = 116, KDiffNet-E, owing to an entry wise parallelizable closed form solution, is on average 2356× faster than the best performing baseline. In Figure S:6(a), we plot the test F1-score for simulated datasets generated using *W* with *p* = 160, representing spatial distances between different 160 regions of the brain. This represents a larger and different set of spatial brain regions. In *p* = 160 case, KDiffNet-E outperforms the best baseline in each case by an average improvement of 67.5%. Including available additional knowledge is clearly useful as JEEK does relatively better than the other baselines, which do not incorporate available additional knowledge. JGLFUSED performs the worst on all cases. Figure S:6(b) shows the computation cost of each method measured in seconds for each case. In all cases, KDiffNet-E has the least computation cost in different cases of varying *n*_*c*_ and *n*_*d*_, as well as with different sparsity of the differential network. KDiffNet-E is on average 3300× faster than the best performing baseline. In Figure S:7(a), we plot the test F1-score for simulated datasets generated using a larger *W* with *p* = 246, representing spatial distances between different 246 regions of the brain. This represents a larger and different set of spatial brain regions. In this case, KDiffNet-E outperforms the best baseline in each case by an average improvement of 66.4% relative to the best performing baseline. Including available additional knowledge is clearly useful as JEEK does relatively better than the other baselines, which do not incorporate available additional knowledge. JGLFUSED performs the worst on all cases. Figure S:7(b) shows the computation cost of each method measured in seconds for each case. In all cases, KDiffNet-E has the least computation cost in different cases of varying *n*_*c*_ and *n*_*d*_, as well as with different sparsity of the differential network. KDiffNet-E is on average 3966× faster than the best performing baseline.

**Figure S:6:**
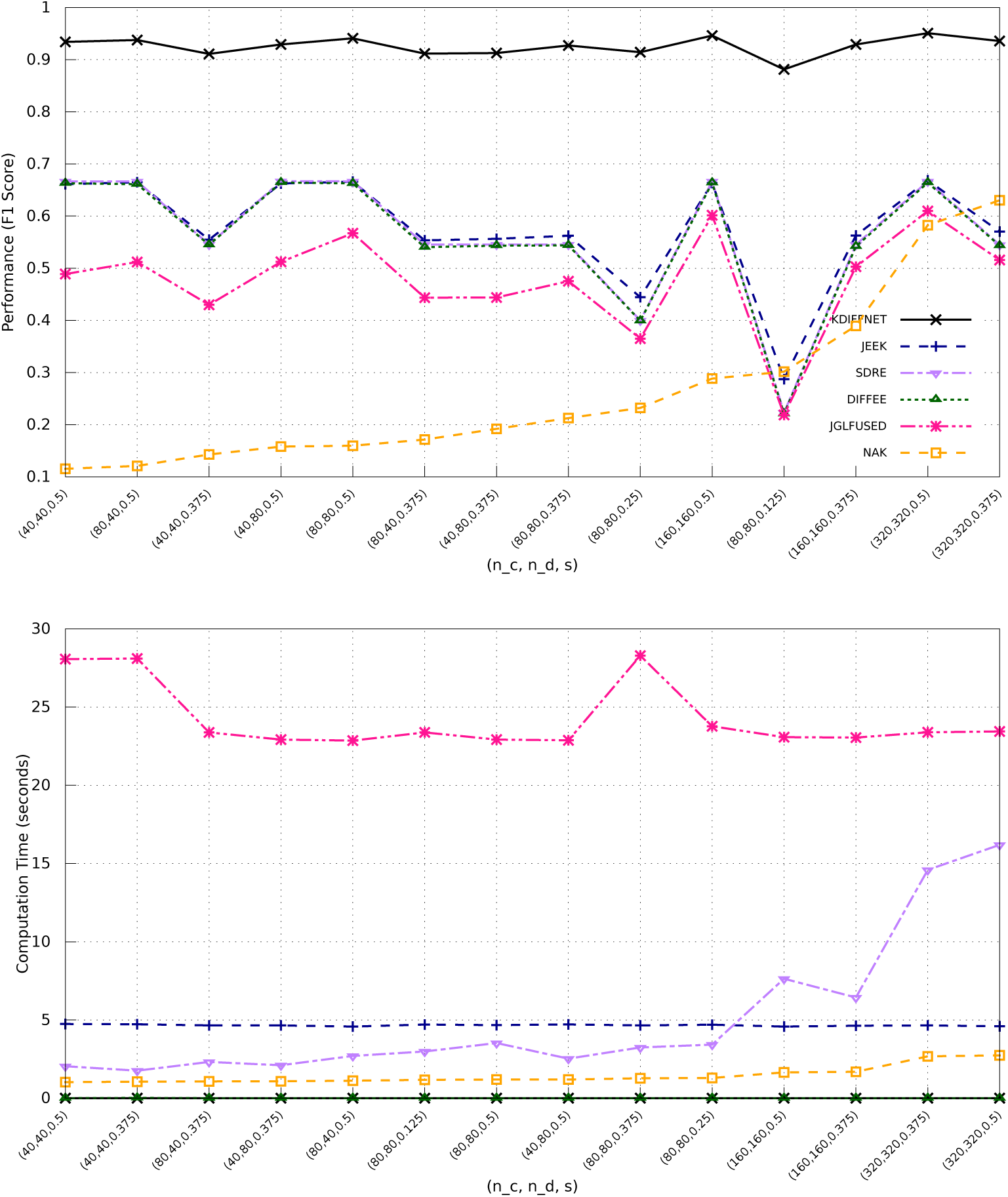
KDiffNet-E Simulation Results for *p* = 160 for different settings of *n*_*c*_, *n*_*d*_ and *s*: (a) The test F1-score and (b) The average computation time (measured in seconds) per *λ*_*n*_ for KDiffNet-E and baseline methods.

**Figure S:7:**
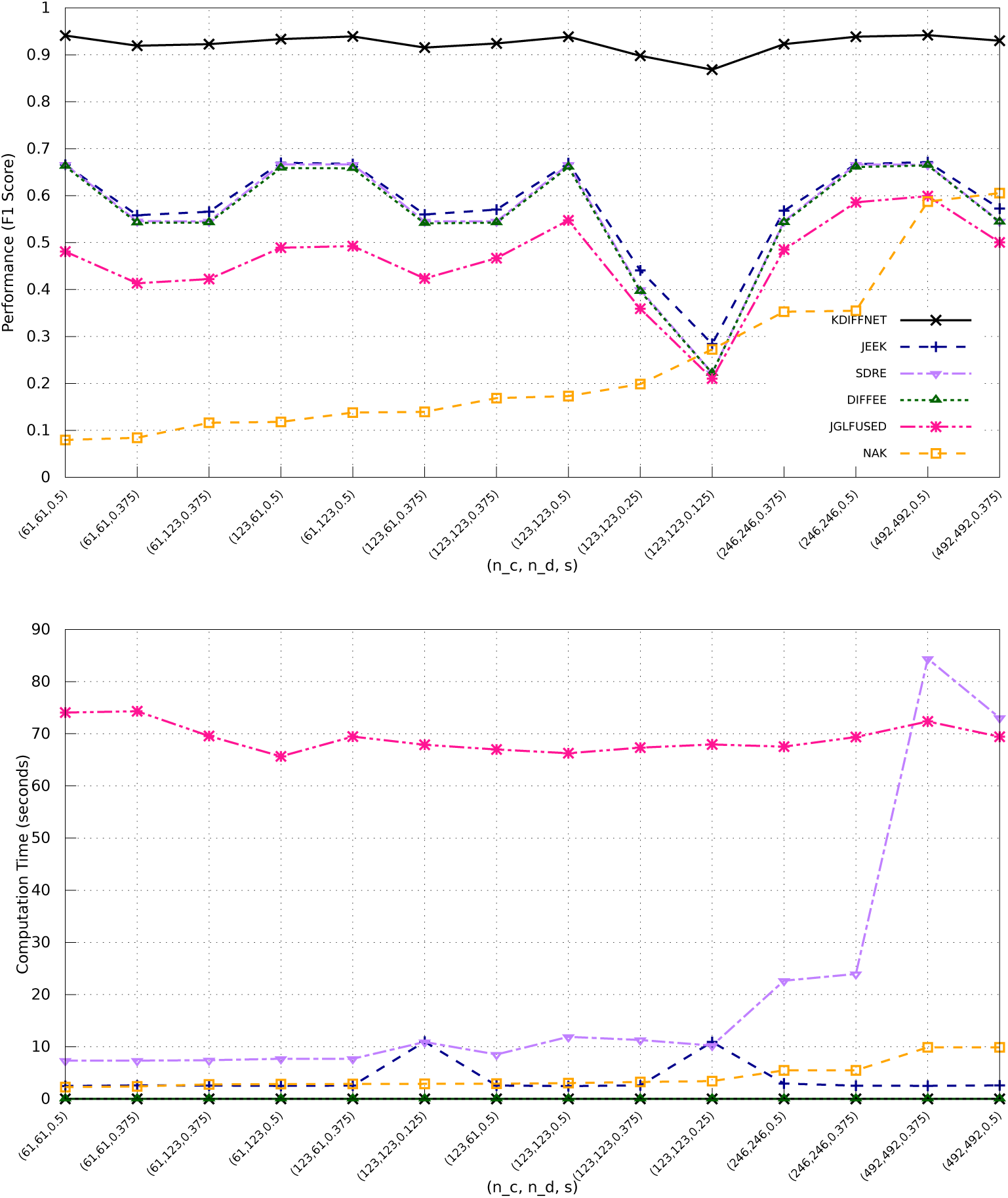
KDiffNet-E Simulation Results for *p* = 246 for different settings of *n*_*c*_, *n*_*d*_ and *s*: (a) The test F1-score and (b) The average computation time (measured in seconds) per *λ*_*n*_ for KDiffNet-E and baseline methods.

###### Node Group Knowledge

We use KDiffNet-G to estimate the differential network with the known groups as extra knowledge. We vary the number of groups *s*_*G*_ and the number of samples *n*_*c*_ and *n*_*d*_ for each case of *p* = {116, 160, 246}. Figure S:8 shows the F1-Score of KDiffNet-G and the baselines for *p* = 116. KDiffNet-G clearly has a large advantage when extra node group knowledge is available. The baselines cannot model such available knowledge.

**Figure S:8:**
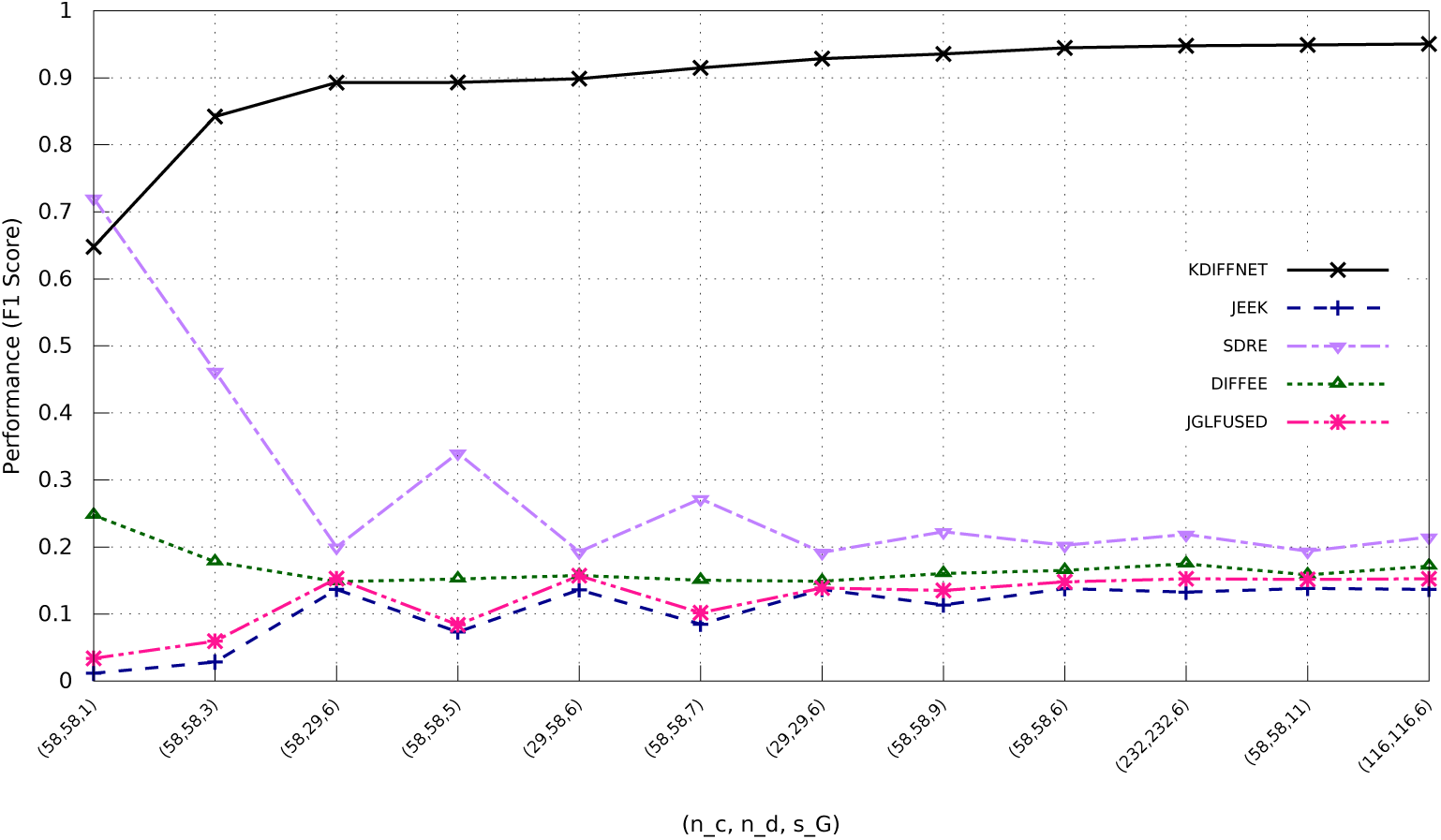
KDiffNet-G Simulation Results for *p* = 246 for different settings of *n*_*c*_, *n*_*d*_ and *s*: (a) The test F1-score and (b) The average computation time (measured in seconds) per *λ*_*n*_ for KDiffNet-E and baseline methods.

###### Varying proportion of known edges

We generate *W*_*E*_ matrices with *p* = 150 using Erdos Renyi Graph [9]. We use the generated graph as prior edge knowledge *W*_*E*_. Additionally, we simulate 15 groups of size 10 as explained in Section S:6.2. We simulate Ω_*c*_ and Ω_*d*_ as explained in Section S:6.2. Figure S:9 presents the performance of KDiffNet-EG, KDiffNet-E and DIFFEE with varying proportion of known edges.

KDiffNet-EG has a higher F1-score than KDiffNet-E as it can additionally incorporate known group information. As expected, with increase in the proportion of known edges, F1-Score improves for both KDiffNet-EG and KDiffNet-E. In contrast DIFFEE cannot make use of additional information and the F1-Score remains the same.

###### Scalability in *p*

To evaluate the scalability of KDiffNet and baselines to large *p*, we also generate larger *W*_*E*_ matrices with *p* = 2000 using Erdos Renyi Graph [9], similar to the aforementioned design. Using the generated graph as prior edge knowledge *W*_*E*_, we design Ω_*c*_ and Ω_*d*_ as explained in Section S:6.2. For the case of both edge and vertex knowledge, we fix the number of groups to 100 of size 10. We evaluate the scalability of KDiffNet-EG and baselines measured in terms of computation cost per *λ*_*n*_.

Figure S:11 shows the computation time cost per *λ*_*n*_ for all methods. Clearly, KDiffNet takes the least time, for large *p* as well.

###### Choice of *λ*_*n*_

For KDiffNet, we show the performance of all the methods as a function of choice of *λ*_*n*_. Figure S:10 shows the True Positive Rate(TPR) and False Positive Rate(FPR) measured by varying *λ*_*n*_ for *p* = 116, *s* = 0.5 and *n*_*c*_ = *n*_*d*_ = *p/*2 under the Data-EG setting. Clearly, KDiffNet-EG achieves the highest Area under Curve (AUC) than all other baseline methods. KDiffNet-EG also outperforms JEEK and NAK that take into account edge knowledge but cannot model the known group knowledge.

**Figure S:9:**
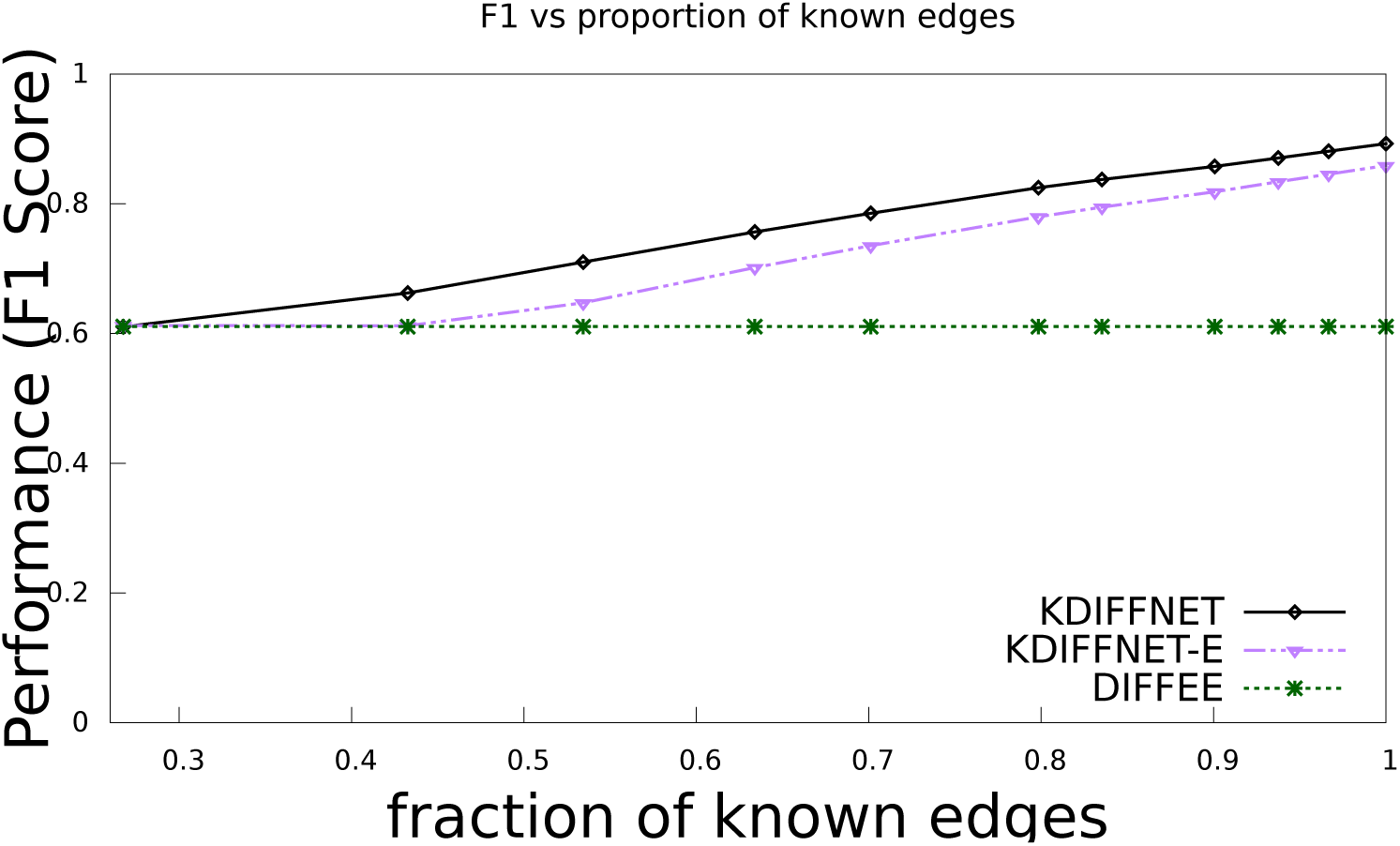
F1-Score of KDiffNet-EG, KDiffNet-E and DIFFEE with varying proportion of known edges.

**Figure S:10:**
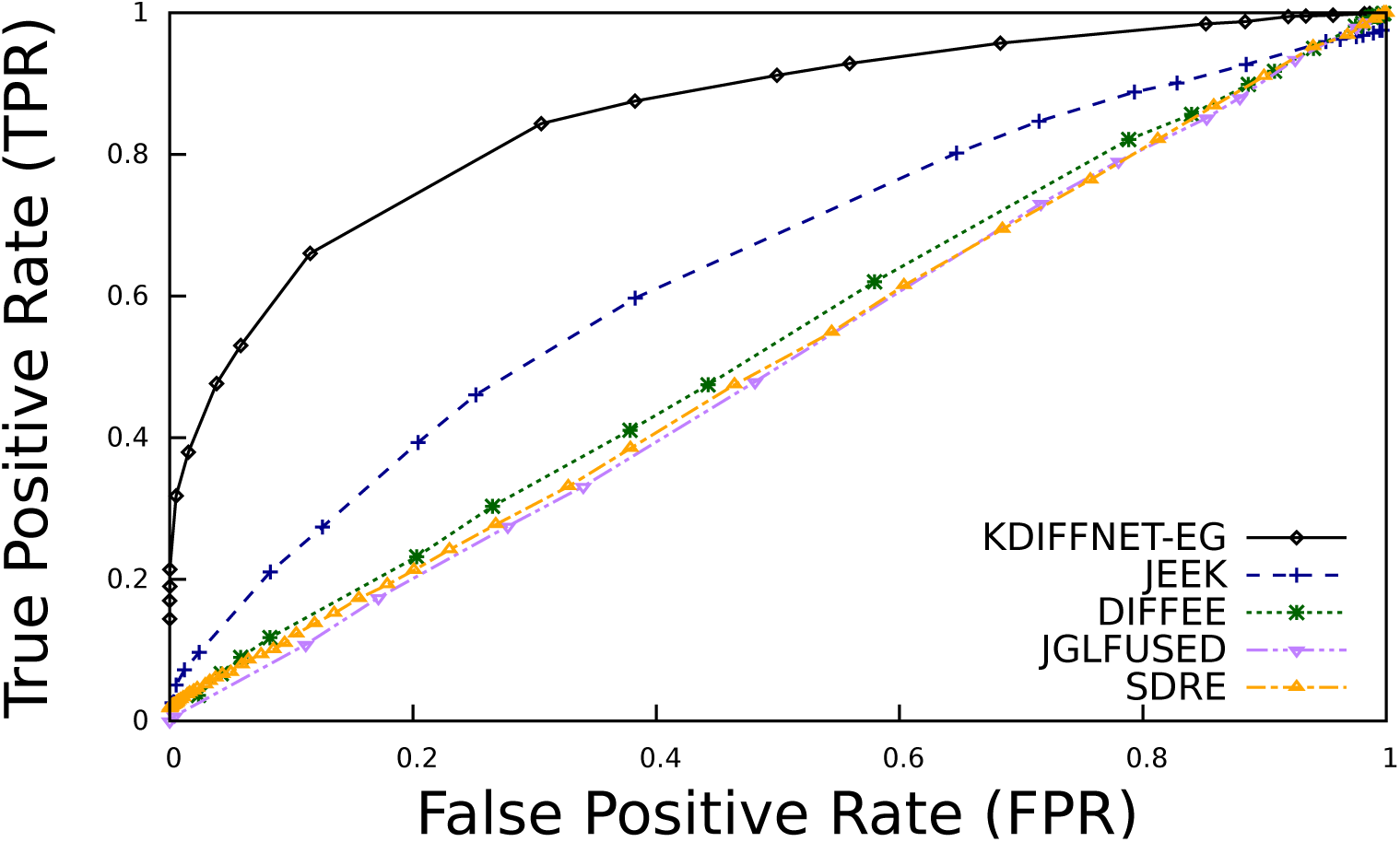
Area Under Curve (AUC) Curves for KDiffNet and baselines at different hyperparameter values *λ*.

**Figure S:11:**
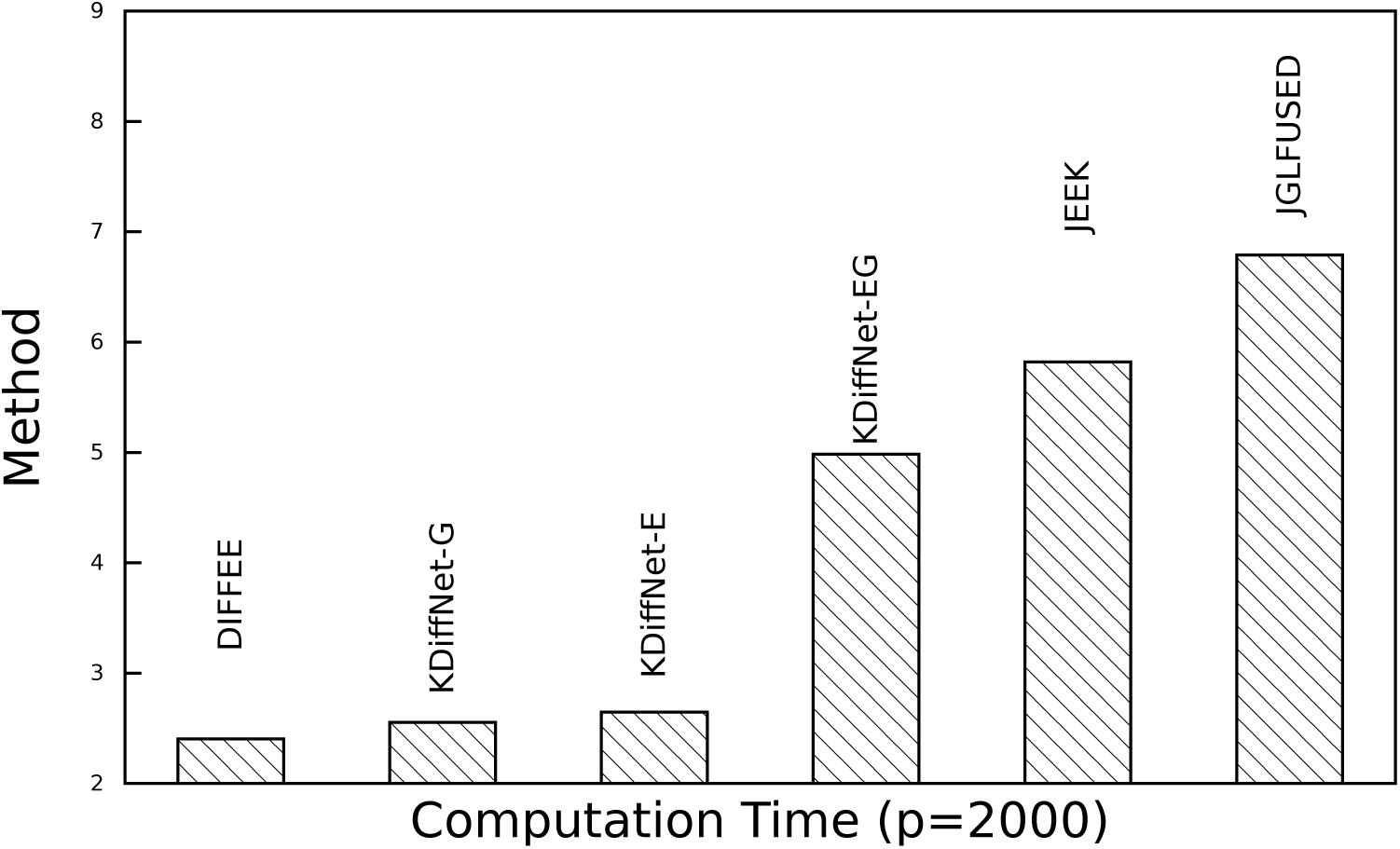
Scalability of KDiffNet : Computation Cost (computation time per *λ*) as a function of *p*.

##### S:6.4 More Experiment: Brain Connectivity Estimation from Real-World fMRI

###### ABIDE Dataset

This data is from the Autism Brain Imaging Data Exchange (ABIDE) [7], a publicly available resting-state fMRI dataset. The ABIDE data aims to understand human brain connectivity and how it reflects neural disorders [19]. The data is retrieved from the Preprocessed Connectomes Project [4], where preprocessing is performed using the Configurable Pipeline for the Analysis of Connectomes (CPAC) [5] without global signal correction or band-pass filtering. After preprocessing with this pipeline, 871 individuals remain (468 diagnosed with autism). Signals for the 160 (number of features *p* = 160) regions of interest (ROIs) in the often-used Dosenbach Atlas [8] are examined. We also include two types of available node groups : one with 40 unique groups of regions belonging to the same functional network and another with 6 node groups about nodes belonging to the same broader anatomical region of the brain.

###### Cross-validation

Classification is performed using the 3-fold cross-validation suggested by the literature [14][20]. We tune over *λ*_*n*_ and pick the best *λ*_*n*_ using cross validation. The subjects are randomly partitioned into three equal sets: a training set, a validation set, and a test set. Each estimator produces 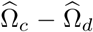 using the training set. Then, these differential networks are used as inputs to Quadratic discriminant analysis (QDA), which is tuned via cross-validation on the validation set. Finally, accuracy is calculated by running QDA on the test set. This classification process aims to assess the ability of an estimator to learn the differential patterns of the connectome structures.

##### S:6.5 Detailed Simulation Results

Table S:2,Table S:3 and Table S:4 present a summary of results for KDiffNet-EG, KDiffNet-E and KDiffNet-G in terms of F1-Score, respectively. We report the average F1-Score(along with standard deviation across the same setting of *n*_*c*_ and *n*_*d*_) across all simulation settings for each *p.* Table S:5,Table S:6 and Table S:7 present a summary of computation time for KDiffNet-EG, KDiffNet-E and KDiffNet-G, respectively. We report the average computation time per *λ*_*n*_ across all simulation settings for each *p*.

---

**Table S:2:**
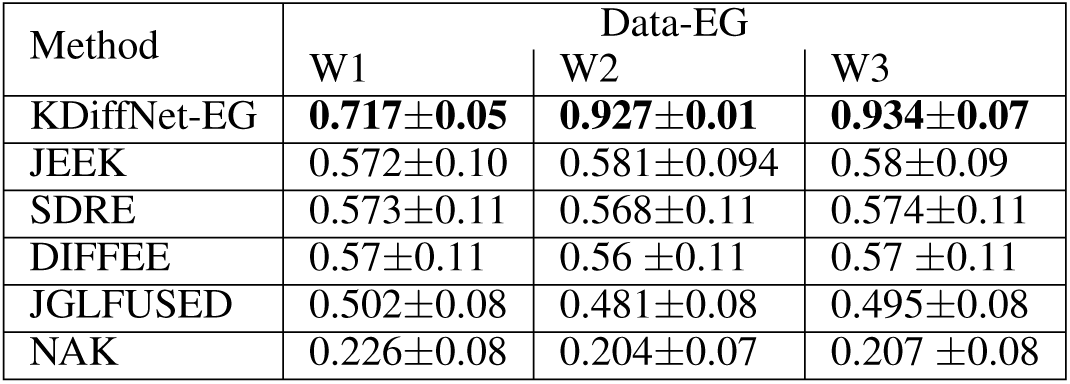
Mean Performance (and standard deviation) of KDiffNet-EG and baselines for multiple data settings.

| Method | Data-EG |  |  |
| --- | --- | --- | --- |
|  | W1 | W2 | W3 |
| KDiffNet-EG | **0.717±0.05** | **0.927±0.01** | **0.934±0.07** |
| JEEK | 0.572±0.10 | 0.581±0.094 | 0.58±0.09 |
| SDRE | 0.573±0.11 | 0.568±0.11 | 0.574±0.11 |
| DIFFEE | 0.57±0.11 | 0.56 ±0.11 | 0.57 ±0.11 |
| JGLFUSED | 0.502±0.08 | 0.481±0.08 | 0.495±0.08 |
| NAK | 0.226±0.08 | 0.204±0.07 | 0.207 ±0.08 |

**Table S:3:** Mean Performance (and standard deviation) KDiffNet-E and baselines for multiple data settings.

| Method | Data-E |  |  |
| --- | --- | --- | --- |
|  | W1 | W2 | W3 |
| KDiffNet-E | **0.687±(0.083)** | **0.924±(0.017)** | **0.926±(0.018)** |
| JEEK | 0.577±(0.09) | 0.581±(0.094) | 0.577±(0.093) |
| SDRE | 0.5643(±0.109) | 0.564(±0.108) | 0.5611(±0.109) |
| DIFFEE | 0.562(±0.109) | 0.5599(±0.109) | 0.564(±0.109) |
| JGLFUSED | 0.484(±0.085) | 0.463(±0.080) | 0.478(±0.081) |
| NAK | 0.274(±0.079) | 0.242(±0.071) | 0.2641(±0.083) |

**Table S:4:**
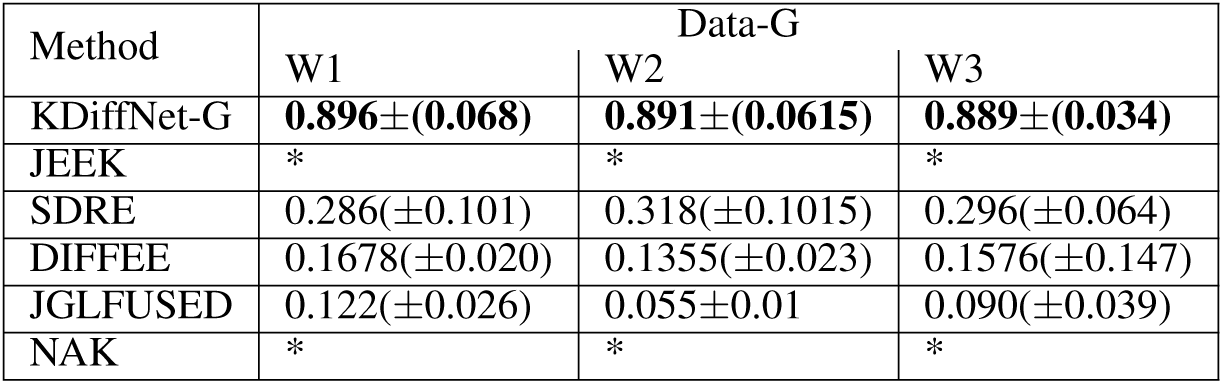
Mean Performance (and standard deviation) of KDiffNet-G and baselines for multiple data settings.

| Method | Data-G |  |  |
| --- | --- | --- | --- |
|  | W1 | W2 | W3 |
| KDiffNet-G | **0.896±(0.068)** | **0.891±(0.0615)** | **0.889±(0.034)** |
| JEEK | * | * | * |
| SDRE | 0.286(±0.101) | 0.318(±0.1015) | 0.296(±0.064) |
| DIFFEE | 0.1678(±0.020) | 0.1355(±0.023) | 0.1576(±0.147) |
| JGLFUSED | 0.122(±0.026) | 0.055±0.01 | 0.090(±0.039) |
| NAK | * | * | * |

**Table S:5:** Mean (and standard deviation) Computation Time (measured in seconds) per *λ*_*n*_ of KDiffNet-EG and baselines for multiple data settings.

| Method | Data-EG |  |  |
| --- | --- | --- | --- |
|  | W1 | W2 | W3 |
| KDiffNet-EG | **0.562±0.077** | 4.591±0.080 | 1.701±0.083 |
| JEEK | 2.460 ±0.020 | 10.998±0.1148 | 4.636±0.023 |
| SDRE | 2.384±0.144 | 27.487±2.5892 | 6.706 ±0.748 |
| DIFFEE | **0.0005 ±0.0002** | 0.004±0.0024 | 0.001 ±0.0002 |
| JGLFUSED | 22.571±3.0053 | 68.128±1.6624 | 24.336±1.450 |
| NAK | 0.780 ±0.0576 | 3.800±0.1518 | 1.432±0.043 |

**Table S:6:** Mean (and standard deviation) Computation Time (measured in seconds) per *λ*_*n*_ of KDiffNet-E and baselines for multiple data settings.

| Method | Data-E |  |  |
| --- | --- | --- | --- |
|  | W1 | W2 | W3 |
| KDiffNet-E | **0.0009±0.0003** | 0.0011±0.0005 | 0.0013±0.0003 |
| JEEK | 2.560±0.1099 | 3.761±1.1835 | 4.661 ±0.1099 |
| SDRE | 1.950±0.264 | 21.024±3.2085 | 5.105±0.2638 |
| DIFFEE | **0.0005 ±0.0002** | 0.004±0.0004 | 0.001 ± 0.0002 |
| JGLFUSED | 10.921±0.3389 | 69.160±2.0849 | 24.252±0.3389 |
| NAK | 0.790±0.061 | 4.235±0.1586 | 1.444±0.0609 |

**Table S:7:** Mean (and standard deviation) Computation Time (measured in seconds) per *λ*_*n*_ of KDiffNet-G and baselines for multiple data settings.

| Method | Data-G |  |  |
| --- | --- | --- | --- |
|  | W1 | W2 | W3 |
| KDiffNet-G | **0.0014±0.0002** | 0.0056±0.0016 | 0.0022±0.0006 |
| JEEK | * | * | * |
| SDRE | 1.685±0.248 | 11.764±1.23 | 4.264±0.566 |
| DIFEE | **0.0009±0.0001** | 0.0035±0.001 | 0.0015±0.0002 |
| JGLFUSED | 22.855±0.593 | 144.47±59.68 | 45.322 ±4.312 |
| NAK | * | * | * |

**Table S:8:** KDiffNet-E : *p* = 160 Relative Performance and speed up with respect to the best performing baseline.

| $n_c$ | $n_d$ | sparsity | Relative F1 Score | Relative Speed up |
| --- | --- | --- | --- | --- |
| 80 | 80 | 0.125 | 3.07 | 2941.13 |
| 40 | 40 | 0.375 | 1.64 | 3376.86 |
| 80 | 40 | 0.375 | 1.65 | 3322.57 |
| 40 | 80 | 0.375 | 1.64 | 4646.60 |
| 80 | 80 | 0.25 | 2.06 | 4696.00 |
| 80 | 80 | 0.375 | 1.65 | 2907.00 |
| 160 | 160 | 0.375 | 1.65 | 2893.00 |
| 40 | 80 | 0.5 | 1.39 | 2102.92 |
| 40 | 40 | 0.5 | 1.40 | 1460.43 |
| 320 | 320 | 0.375 | 1.64 | 2584.67 |
| 80 | 40 | 0.5 | 1.41 | 2698.00 |
| 80 | 80 | 0.5 | 1.41 | 2935.75 |
| 160 | 160 | 0.5 | 1.42 | 6359.17 |
| 320 | 320 | 0.5 | 1.42 | 3285.71 |

**Table S:9:**
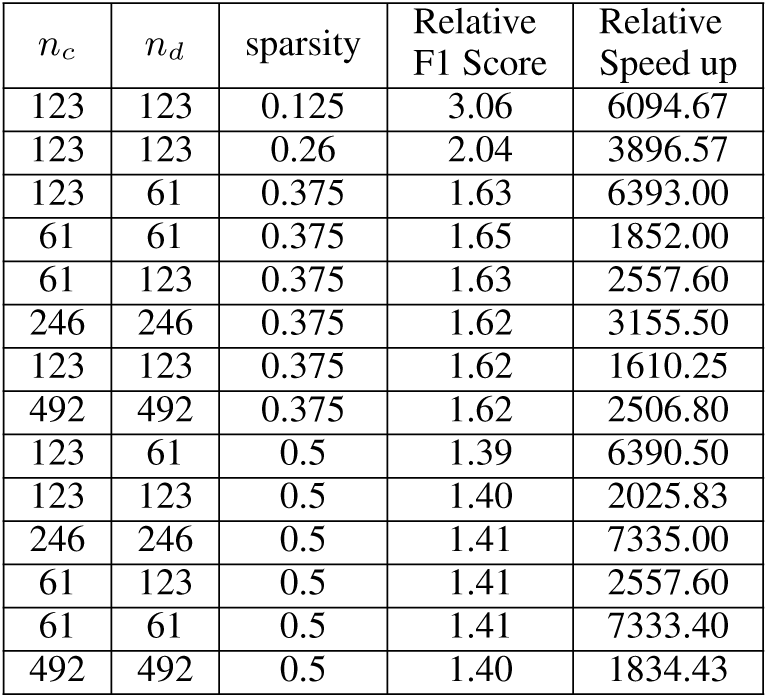
KDiffNet-E : *p* = 246 Relative Performance and speed up with respect to the best performing baseline.

| $n_c$ | $n_d$ | sparsity | Relative F1 Score | Relative Speed up |
| --- | --- | --- | --- | --- |
| 123 | 123 | 0.125 | 3.06 | 6094.67 |
| 123 | 123 | 0.26 | 2.04 | 3896.57 |
| 123 | 61 | 0.375 | 1.63 | 6393.00 |
| 61 | 61 | 0.375 | 1.65 | 1852.00 |
| 61 | 123 | 0.375 | 1.63 | 2557.60 |
| 246 | 246 | 0.375 | 1.62 | 3155.50 |
| 123 | 123 | 0.375 | 1.62 | 1610.25 |
| 492 | 492 | 0.375 | 1.62 | 2506.80 |
| 123 | 61 | 0.5 | 1.39 | 6390.50 |
| 123 | 123 | 0.5 | 1.40 | 2025.83 |
| 246 | 246 | 0.5 | 1.41 | 7335.00 |
| 61 | 123 | 0.5 | 1.41 | 2557.60 |
| 61 | 61 | 0.5 | 1.41 | 7333.40 |
| 492 | 492 | 0.5 | 1.40 | 1834.43 |

## Footnotes

1 We put details of theoretical proofs, details of how we generate simulation datasets and concrete performance figures in the appendix. Notations with “S:” as the prefix indicate the corresponding contents are in the appendix.

2 For instance, on samples from a controlled drug study ‘c’ may represent the ‘control’ group and ‘d’ may represent the ‘drug-treating’ group. Using which of the two sample sets as ‘c’ set (or ‘d’ set) does not affect the computational cost and does not influence the statistical convergence rates.

3 We use the same range to tune *λ*_1_ for SDRE and *λ* _2_ for JGLFUSED. We use *λ* _1_ = 0.0001(a small value) for JGLFUSED to ensure only the differential network is sparse. Tuning NAK is done by the package itself.

4 We cannot compare to NAK and SDRE because they do not provide precision matrix required for QDA

1 http://allmodelsarewrong.net/kliep_sparse/demo_sparse.html

2 This indicates for some positive constant 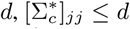 and 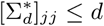 for all diagonal entries. Moreover, if *q* = 0, then this condition reduces to 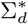 and 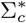 being sparse.

3 The machine that we use for experiments is an Intel Core i7 CPU with a 16 GB memory.

